# A Cold-Responsive Mitochondrial Transporter Stimulates Brown Adipose Tissue Thermogenesis

**DOI:** 10.64898/2026.06.30.735039

**Authors:** Yin Long, Xu Yang, Jiayu Zhou, Jieyuan Xue, Kaimin Wu, Fang Chen, Wenyan Li, Hongyong Song, Kaihua Zhang, Xu-Yun Zhao

## Abstract

Metabolites are emerging as signaling molecules that mediate cellular function, extending beyond their well-established roles in metabolic pathways. Members of the solute carrier (SLC) family mediate metabolite transport across cellular compartments, raising the possibility that these proteins may sense environmental stimuli and regulate cellular biological processes by triggering signaling cascades linked to metabolite transport. This study investigated the response of the SLC25A family, a unique set of inner mitochondrial membrane-localized transporters, to cold as an environmental stimulus in mediating metabolic reprogramming; and whether this reprogramming, driven by the metabolites transported by SLC25A proteins, subsequently promotes the activation of thermogenesis in brown adipocytes. After screening members of the SLC25A family for their responsiveness to cold stimuli and brown adipose tissue (BAT) activation, we found that Slc25a34 was robustly induced under these conditions. We further demonstrated that Slc25a34 mediates the transport of adenosine monophosphate (AMP), derived from *de novo* glucose synthesis, from mitochondria to the cytosol. This transport potentiates AMP-activated protein kinase (AMPK) signaling and glycolytic flux in brown adipocytes, both of which facilitate BAT thermogenesis during cold exposure. Intriguingly, cold exposure directly promoted the activation of peroxisome proliferator-activated receptor gamma (PPARγ), which transcriptionally upregulated *Slc25a34* expression. More importantly, genetic ablation of *Slc25a34* impaired BAT thermogenesis. Thus, our study reveals a novel cold-induced metabolite-sensing pathway, where Slc25a34-mediated AMP transport between mitochondria and the cytosol serves as a critical signal for activating BAT thermogenesis. These findings provide compelling evidence that metabolite transport across cellular compartments acts as a key driver of cellular physiology, thereby offering novel insights into metabolite-based therapeutic strategies for metabolic diseases.

**Highlights:**

- Slc25a34 is cold-responsive and transcriptionally regulated by PPARγ.
- Slc25a34 functions specifically to mediate the mitochondrial-to-cytosolic transport of AMP in brown adipocytes.
- Mitochondrially sequestered *de novo* synthesized AMP acts as a signaling reservoir, and its Slc25a34-mediated efflux to the cytosol activates AMPK and glycolysis, supporting BAT thermogenesis.

## Introduction

In recent years, the concept of metabolites as signaling molecules in regulating cellular homeostasis has garnered significant attention ^1, 2^. Traditionally dismissed as mere intermediates in metabolic pathways, metabolites are now widely recognized for their critical roles in orchestrating cellular signaling and regulatory processes ^3^. This paradigm shift has spurred growing interest in unraveling the interplay between metabolic states and cellular regulatory mechanisms ^4, 5^. Emerging research indicates that a cell’s physiological state reflects a dynamic interplay between its regulatory systems and intermediary metabolism, a reciprocity where regulatory networks not only shape metabolic dynamics but are themselves modulated by feedback from metabolic states ^6^. This bidirectional crosstalk highlights metabolites as central hubs linking metabolism to function, yet many of its underlying mechanisms remain to be fully elucidated.

Metabolic reprogramming mediates cellular signaling through diverse metabolite-sensing mechanisms, including direct binding to sensors, interaction with G protein-coupled receptors (GPCRs) ^7^, and regulation of protein modifications ^8^. However, while these mechanisms have been extensively characterized, the extent to which metabolite transport across subcellular compartments, particularly between mitochondria and the cytosol, drives signaling cascades remain poorly understood.

Mitochondria, often termed the “powerhouses of the cell,” play a pivotal role in cellular metabolism by regulating metabolite flux across their inner membrane, a process essential for sustaining core metabolic reactions such as the tricarboxylic acid (TCA) cycle, oxidative phosphorylation, and fatty acid oxidation ^9^. Central to this function is the solute carrier 25 (SLC25) family of transporters, which mediate the exchange of metabolites, nucleotides, and cofactors across the mitochondrial inner membrane, thereby linking mitochondrial metabolism to cytosolic processes ^10^. For instance, SLC25A44 facilitates the transport of branched-chain amino acids (BCAAs) into mitochondria, a process critical for energy homeostasis and thermogenesis ^11, 12^, while SLC25A33 and SLC25A36 mediate pyrimidine nucleotide transport, supporting DNA and RNA synthesis ^13^. Dysregulation or mutations in these transporters disrupt metabolic balance, contributing to diseases ranging from metabolic syndrome to neurodegeneration ^14^, underscoring the importance of mitochondrial metabolite transport in health and disease.

Against this backdrop, cold-induced thermogenesis in brown adipose tissue (BAT) represents a robust physiological model to study the crosstalk between metabolic processes and the regulation of gene expression. BAT is uniquely specialized to dissipate energy as heat either via uncoupling protein 1 (UCP1) or through several UCP1-independent mechanisms, a process critical for maintaining core body temperature during cold exposure and regulating systemic energy expenditure^15^. Cold stimuli activate BAT through well-characterized pathways, including the β-adrenergic receptor (β-AR) signaling cascade, which triggers downstream events such as cyclic AMP (cAMP) production and protein kinase A (PKA) activation^16^. Additionally, activation of the AMP-activated protein kinase (AMPK) pathway is pivotal for enhancing thermogenic gene expression in adipose tissue, thereby increasing energy expenditure and improving metabolic health ^17, 18^. Pharmacological activation of AMPK has even been demonstrated to confer protection against diet-induced obesity via the induction of thermogenic fat activation ^19^. Beyond signaling, transcriptional networks, including the key regulator peroxisome proliferator-activated receptor gamma (PPARγ), orchestrate thermogenic gene expression, linking UCP1 activation to BAT’s overall thermogenic capacity ^20^. Despite these advances, how cold exposure directly modulates metabolic homeostasis of BAT and how such metabolic reprogramming influences thermogenic capacity remain elusive.

To address this gap, we set out to identify mitochondrial transporters that bridge metabolic reprogramming and thermogenic signaling in BAT. We screened the RNA expression profiles of SLC25A family genes in several cold-induced thermogenesis mouse models. This approach led to the identification of Slc25a34 as a gene whose expression is robustly induced by cold exposure and β-adrenergic receptor (β-AR) stimulation in BAT. Using *Slc25a34* knockout mice, we demonstrated that Slc25a34 is indispensable for adaptive thermogenesis in BAT. Mechanistically, *Slc25a34*-deficient brown adipocytes exhibited disrupted purine metabolism due to impaired translocation of adenosine monophosphate (AMP) from mitochondria to the cytosol. This defect hindered cytosolic AMPK phosphorylation and glycolytic activity, ultimately impairing BAT thermogenic function.

In summary, our study uncovers a novel cold-induced metabolic reprogramming mechanism in BAT, where Slc25a34-mediated mitochondrial-cytosolic AMP transport serves as a critical signaling node activating thermogenesis. These findings underscore that metabolite shuttling between subcellular compartments acts as a key modulator of cellular signaling that augments adipose thermogenesis, thereby offering critical mechanistic insights for designing innovative anti-obesity therapeutics targeting adipose thermogenic activation.

## Results

### Slc25a34 is robustly induced in brown adipocytes in response to cold stimulation via the β3-adrenergic pathway

To explore whether transporters of the SLC25A family can sense environmental stimuli, and subsequently rewire metabolic homeostasis to govern physiological processes, we employed cold exposure as a relevant physiological model. We first profiled the expression of SLC25A family genes in mouse BAT under distinct experimental conditions including two cold exposure paradigms, chronic cold (CC) and acute cold (AC), with room temperature (RT) as the control, and direct treatment with the β3-adrenergic agonist CL316,243 (CL), with saline injected mice as the control. Quantitative PCR (qPCR) analysis demonstrated that among all SLC25A family members, Slc25a34 exhibited the most dramatic upregulation in BAT upon CC, AC, or CL treatment relative to their controls (Figure 1A). Notably, CL administration also markedly increased *Slc25a34* expression in inguinal white adipose tissue (iWAT), whereas acute and chronic cold exposure had only mild effects in this depot (Figure S1A). To evaluate the dynamic induction of *Slc25a34* by cold, we performed a temperature reversal experiment. We observed that following CC treatment, *Slc25a34* mRNA levels in BAT were elevated but returned to near-basal levels when mice were re-exposed to room temperature (Figure S1B).

**Figure 1.**
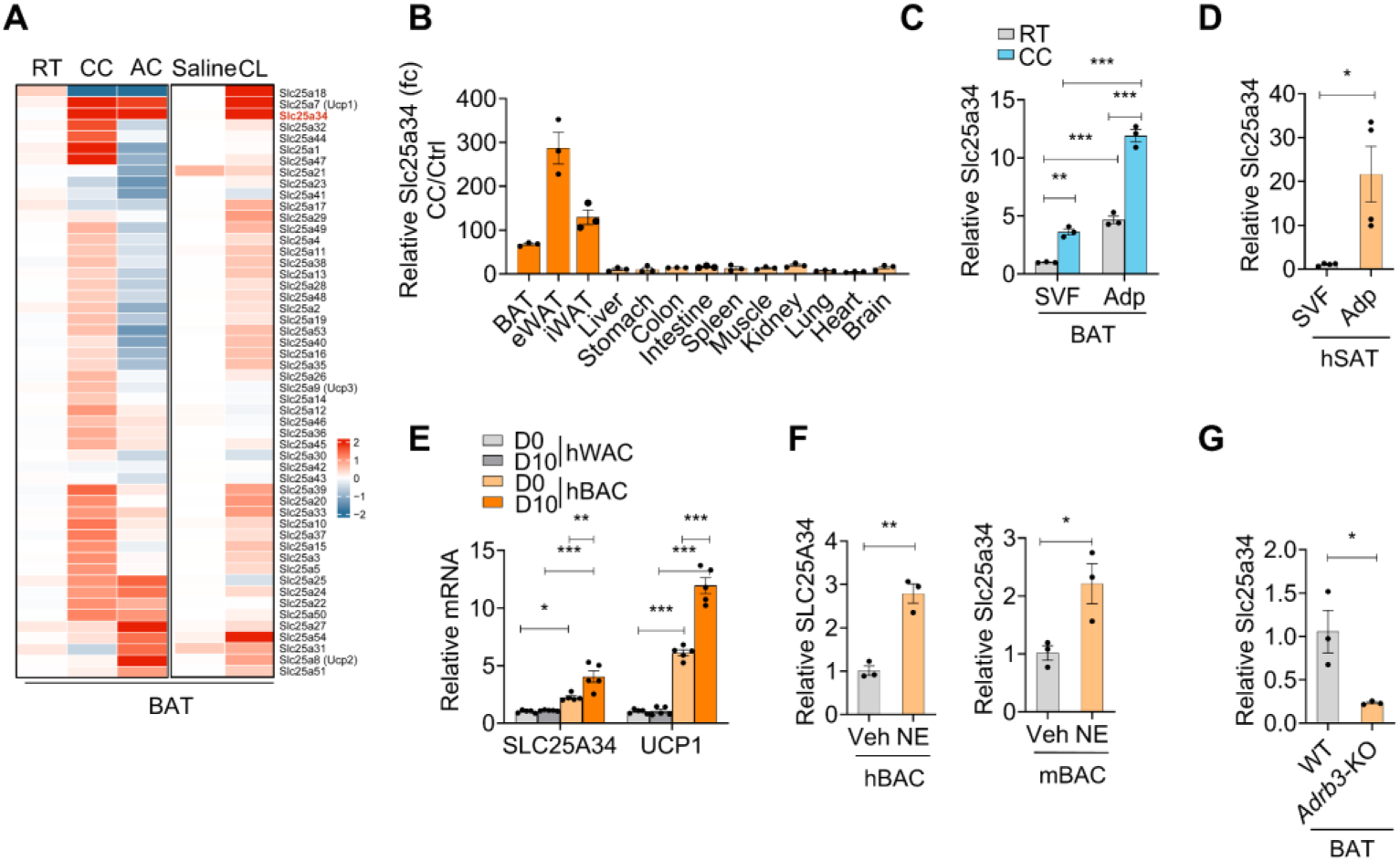
SLC25A34 is highly induced in brown adipocytes by cold-induced β3-adrenergic signaling. (**A**) QPCR analysis of the mRNA expression of SLC25A transporter family members in BAT from wild type (WT) mice kept in room temperature (RT, n = 4), exposed to chronic cold (CC, n = 3), exposed to acute cold (AC, n = 4) and injected with CL316,243 (CL, 1 mg/kg, n = 4) or saline (Sal, n = 4) for 7 consecutive days. (**B**) The mRNA level of Slc25a34 in various tissues from WT mice maintained in room temperature and exposed to chronic cold for 7 days (n = 3) were determined via qPCR; The fold change was calculated by dividing the mRNA level of Slc25a34 in mice exposed to chronic cold by that housed in room temperature. (**C**) The mRNA expression of Slc25a34 in stromal vascular fraction (SVF) and adipocyte fraction (Adp) of brown adipose tissue (BAT) from WT mouse housed in room temperature (RT, n = 3) and exposed to chronic cold (CC, n = 3) were determined via qPCR. (**D**) The mRNA of SLC25A34 in SVF and Adp of human subcutaneous adipose tissue (hSAT, n = 4) were determined via qPCR. (**E**) mRNA expression of SLC25A34 in the human white adipocyte (hWAC) and brown adipocyte (hBAC) before (day 0) and after differentiation (day 10) (n = 5) were determined via qPCR. (**F**) The mRNA expression of Slc25a34 in differentiated human BAC (left) and mouse BAC (right) treated with vehicle (Veh) or norepinephrine (NE, 1 µM, 6 h) (n = 3) were analyzed by qPCR. (**G**) The mRNA expression of Slc25a34 in the BAT of WT and *Adrb3*-KO mice (n = 3) were determined via qPCR. Data are presented as mean ± SEM. Statistical significance was determined by unpaired two-tailed Student’s t-test in (D), (F) and (G); two-way ANOVA multiple comparison test in (C) and (E). *P <0.05, **P <0.01, ***P <0.001. ns: not significant.

To further elucidate the adipose tissue-specific induction of *Slc25a34* upon cold stimulation, we quantified its expression across major organs. Chronic cold exposure induced a more robust upregulation of *Slc25a34* expression in BAT, iWAT, and epididymal white adipose tissue (eWAT) relative to other tissue types, confirming the adipose-selective regulation of this gene (Figure 1B).

To further identify the cellular source of cold-responsive *Slc25a34* expression, we separated adipose tissue into stromal vascular fractions (SVF) and adipocyte fractions and analyzed *Slc25a34* expression in these components. We revealed that Slc25a34 mRNA levels were higher in adipocytes of BAT than in SVF, and chronic cold treatment further induced its expression in adipocytes (Figure 1C). Consistently, SLC25A34 was also upregulated in the adipocyte fraction compared to SVF in human subcutaneous adipose tissue (hSAT) further confirming this adipocyte-enriched expression (Figure 1D).

Beyond cold responsiveness, *Slc25a34* expression increased progressively during adipocyte differentiation, with a more robust induction in brown adipogenesis from brown adipocyte precursors (BAC) relative to white adipogenesis from C3H10T1/2 (10T1/2) cells (Figure S1C). Furthermore, we measured SLC25A34 expression in human brown adipocytes (hBAC) and white adipocytes (hWAC), isolated from human brown and white fat depots respectively, both before and after differentiation. Consistently, SLC25A34 expression was significantly upregulated during human brown adipocyte differentiation, with higher levels observed in both precursor and mature brown adipocytes relative to white adipocyte counterparts. (Figure 1E).

To further define the cold-induced stimuli driving Slc25a34 expression, we treated differentiated BAC with norepinephrine (NE), a catecholamine that activates adrenergic receptors. NE treatment directly upregulated *Slc25a34* expression in both human and mouse BAC (Figure 1F). More importantly, knockout of the β3-adrenergic receptor (β3-AR) markedly reduced *Slc25a34* expression in BAT, indicating that Slc25a34 may be directly induced by NE-driven β3-adrenergic signaling (Figure 1G).

### SLC25A34 mediates mitochondrial AMP transportation

To dissect the function of SLC25A34 in BAC metabolic reprogramming and its consequent effects on thermogenesis and metabolic health, we generated *Slc25a34*-knockout (KO) mice and established immortalized wild-type (WT) and *Slc25a34*-KO BAC lines. *Slc25a34* knockout efficiency was confirmed in major tissues including BAT, and KO BAC lines (Figure S1D-F), followed by analysis of global metabolic alterations in whole-cell lysates and purified mitochondria. Our metabolomic analysis revealed pronounced dysregulation of nucleotide metabolism in *Slc25a34*-KO BAC, characterized by elevated mitochondrial levels of nucleotide mono-, di- and triphosphates (NMPs, NDPs and NTPs) and increased whole-cell NMPs (Figure 2A-B). Conversely, whole-cell levels of nucleotide di- and triphosphates (NDPs and NTPs) were significantly reduced in *Slc25a34*-KO BAC compared to WT control, indicating Slc25a34 mediates NMPs metabolism in mitochondria which influences whole cell NDPs and NTPs production.

**Figure 2.**
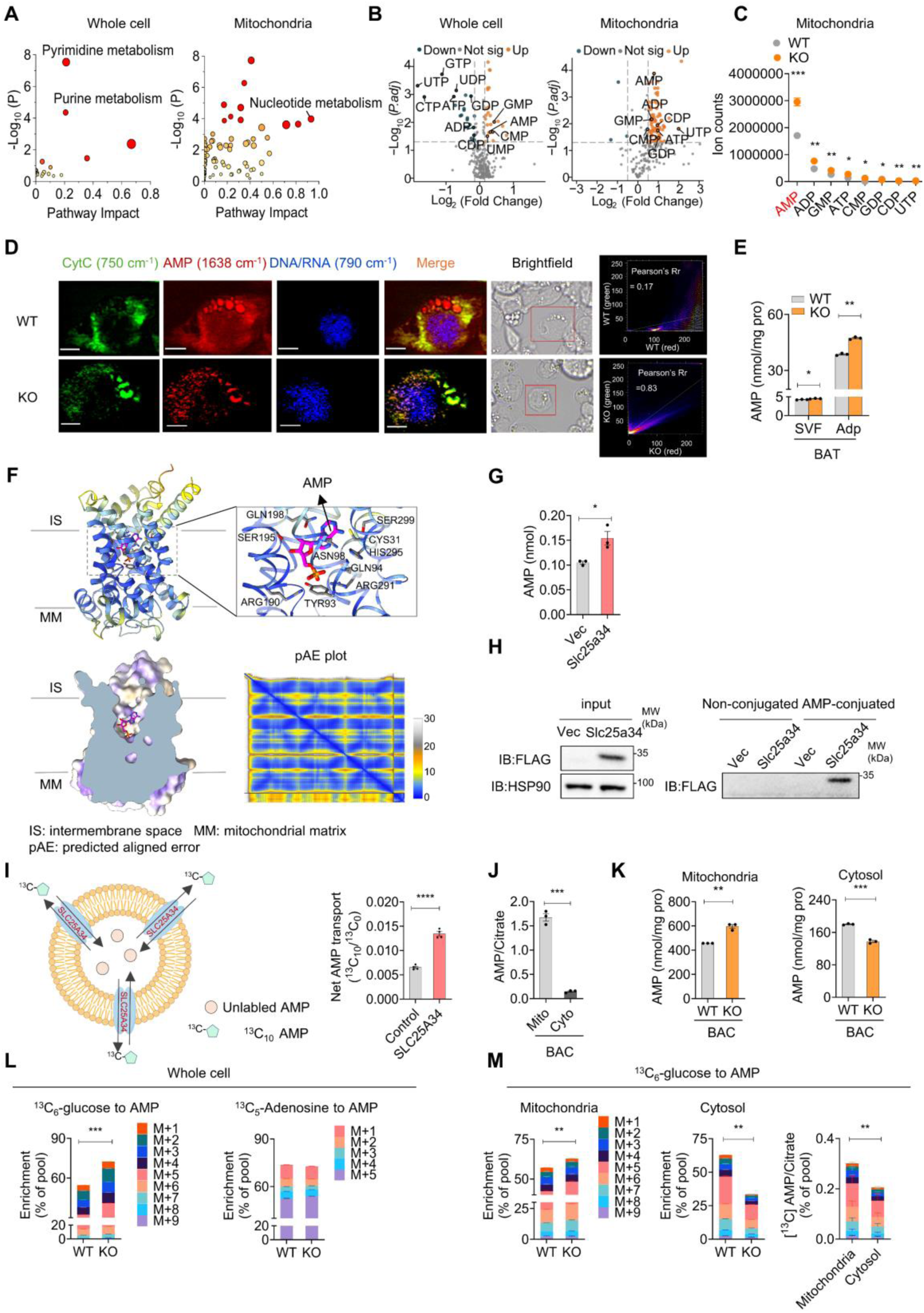
SLC25A34 mediates the transport of AMP from mitochondria to the cytosol. (**A-B**) Pathway enrichment (A) and Volcano plot (B) analysis of differentially expressed metabolites in whole cells (left) and mitochondria (right) between wild type (WT) and *Slc25a34*-KO brown adipocytes (BAC) after differentiation (n = 3). (**C**) The relative abundance of nucleotides in mitochondria from differentiated WT and *Slc25a34*-KO BAC (n = 3). (**D**) Raman images showed the colocalization of mitochondria and AMP in differentiated WT and *Slc25a34*-KO BAC. The Raman peak at 750 cm^-1^ corresponds to CytC (green), indicating mitochondrial distribution. The Raman peak at 1638 cm^-1^ was assigned to AMP (red), whereas the Raman peak at 790 cm^-1^ was used to identify nucleotides in the cell nucleus (blue). Brightfield images are shown for reference. Scale bar=7 µm. The colocalization of AMP and CytC signals was quantified using pearson’s correlation coefficient (Pearson’s Rr). (**E**) AMP concentration was measured in stromal vascular fraction (SVF) and adipocyte fraction (Adp) of BAT from WT and *Slc25a34*-KO mice (n = 3). (**F**) AlphaFold-predicted structure of SLC25A34 colored by pLDDT (left) and a zoom-in view of the AMP-binding pocket (right) showing key residues (grey stick) of SLC25A34 interacting with the ligand (magenta carbon) in the top row. Cutaway side-view of the hydrophobicity surface is shown to present the binding pose of AMP in the pocket (left) and pAE is plotted to highlight the confidence in the relative position of the protein and the ligand (right) in the bottom row. A blue tile indicates a good prediction, whereas a grey tile corresponds to a poor prediction (see color key). (**G**) AMP content in immunoprecipitated anti-FLAG antibody conjugated beads from HEK 293T cells transfected with vector (Vec) or Flag-tagged Slc25a34 (n = 3). (**H**) HEK 293T cells transfected with Vec or Flag-Slc25a34 were subjected to co-immunoprecipitation (Co-IP) using AMP-conjugated beads. The immunoprecipitated SLC25A34 was examined by western blot. (**I**) Schematic of AMP transport assays (left) and SLC25A34 protein was reconstituted into proteoliposomes. ^13^C10-labeled AMP was incubated with empty proteoliposomes or those expressing SLC25A34 for 20 min, detected by LC-MS (right) (n = 4). (**J**) The ratio of AMP to citrate levels in mitochondria and cytosol of differentiated mouse BAC (n = 3). (**K**) The AMP level in mitochondria (left) and cytosol (right) in differentiated WT and *Slc25a34*-KO mouse BAC (n = 3). (**L**) Mass isotopologue distribution analysis of AMP in whole cell of differentiated WT and *Slc25a34*-KO mouse BAC following 24-hour incubation with ^13^C6-glucose (left) and ^13^C5-adenosine (right). (**M**) Mass isotopologue distribution analysis of AMP in mitochondria (left) and cytosol (right) of differentiated WT and *Slc25a34*-KO mouse BAC following 24-hour incubation with ^13^C6-glucose. Data are presented as mean ± SEM. Statistical significance was determined by unpaired two-tailed Student’s t-test in (C), (E),, (I), (J), (K), (L) and (M). *P <0.05, **P <0.01, ***P <0.001. ns: not significant.

To identify the most significantly affected nucleotides in mitochondria by Slc25a34, we ranked mitochondrial nucleotides by their absolute abundance in both WT and *Slc25a34*-KO BAC. This analysis revealed that AMP showed the most striking accumulation in mitochondria of *Slc25a34*-KO BAC (Figure 2C). To characterize the subcellular distribution of AMP in BAC, we employed Raman microscopy to visualize endogenous AMP and cytochrome C in intact cells. Strikingly, AMP signals showed extensive colocalization with cytochrome C in *Slc25a34*-knockout (KO) BAC (Figure 2D), revealing that *Slc25a34* deficiency drives AMP accumulation within mitochondria. We next validated this phenotype via FITC labeling of AMP. In *Slc25a34*-KO BAC, the FITC signal also exhibited a condensed pattern with clear overlap with MitoTracker, whereas in WT BAC, the FITC signal was diffuse and showed distinct segregation from the mitochondrial MitoTracker signal (Figure S2A). Together, these two lines of evidence establish that ablation of Slc25a34 shifts AMP localization toward the mitochondrial compartment. To further confirm that AMP specifically accumulates in adipocytes within *Slc25a34*-KO BAT, we measured AMP, ADP and ATP levels in both the SVF and adipocyte fraction of BAT. We found that AMP levels were significantly higher in both the SVF and adipocyte fraction of *Slc25a34*-KO BAT compared to WT. Notably, AMP was more accumulated in the adipocyte fraction of *Slc25a34*-KO BAT, whereas no such accumulation was observed for ADP or ATP (Figure 2E and S2B-C), suggesting SLC25A34 is critical in regulating AMP homeostasis especially in adipocytes.

To date, neither a physiological substrate for SLC25A34 nor a dedicated mitochondrial transporter for AMP has been identified or characterized. Using AlphaFold3^21^, we predicted the binding poses of AMP in SLC25A34. The confident high-quality prediction positions AMP within the central path of SLC25A34. The structural model further reveals polar interactions between AMP and the surrounding helical bundles, which reinforces a potential role of SLC25A34 in AMP transport (Figure 2F). Co-immunoprecipitation (CoIP) specifically captured SLC25A34, and subsequent AMP quantification showed significantly higher AMP levels in the immunoprecipitated complex from Slc25a34-overexpressing cells compared to control cells. This further validates the interaction between SLC25A34 and AMP. Notably, ADP and ATP were not enriched in these complexes, which further confirms the specificity of SLC25A34 for AMP binding (Figure 2G and Figure S2D). Conversely, AMP conjugated agarose beads were also able to pull down SLC25A34 but not other SLC25A family members such as SLC25A20 (Figure 2H and Figure S2E). Furthermore, mutation of the predicted AMP-binding residues abrogated the binding between SLC25A34 and AMP, further validating the specific interaction of SLC25A34 with AMP (Figure S2E). To elucidate whether SLC25A34 interacts with AMP and transports it between cellular compartments, we performed a proteoliposome-based transport assay. We showed that *in vitro* reconstituted SLC25A34-containing proteoliposomes are capable of importing ^13^C10- and FITC-labeled AMP into the liposomes, as indicated by LC-MS and fluorescence accumulation (Figure 2I and S2F). Collectively, these findings demonstrate that SLC25A34 mediates AMP transport between the mitochondria and the cytosol.

AMP is a key cytosolic nucleotide metabolite that serves as a precursor for ADP and ATP synthesis, yet the presence and functional role of mitochondrial AMP remain poorly characterized. To address this, we next mapped the subcellular distribution of AMP between mitochondria and the cytosol in brown adipocytes. Normalizing to subcellular citrate levels, a metabolite with known mitochondrial-cytosolic shuttling^22^, we found that mitochondrial AMP levels were significantly higher than those in the cytosol (Figure 2J, Figure S2G), whereas ATP showed the opposite pattern, being more abundant in the cytosol (Figure S2H). To investigate the direction of SLC25A34-mediated AMP transport, we measured AMP levels in both compartments of WT and *Slc25a34*-KO BAC. In *Slc25a34*-KO BAC, AMP accumulated in mitochondria and decreased in the cytosol (Figure 2K). Parallel observations were recapitulated in *Slc25a34*-deficient BAC from both Raman imaging of endogenous AMP and staining with FITC-conjugated AMP (Figure 2D and S2A). These findings indicate that SLC25A34 mediates AMP efflux from mitochondria.

In mammalian cells, AMP is produced via two primary pathways, *de novo* purine synthesis and the purine salvage pathway, which utilize glucose and adenosine as substrates, respectively ^23, 24^. To investigate the origin of mitochondrial AMP transported by SLC25A34, we incubated differentiated WT and *Slc25a34*-KO BAC with isotopically labeled tracers, ^13^C6-glucose and ^13^C5-adenosine, and tracing their derived nucleotide metabolites. Results revealed that AMP containing 1 to 9 carbons derived from ^13^C6-glucose (designated as M+1 to M+9) were significantly increased in *Slc25a34*-KO BAC. In contrast, no significant differences were observed in AMP containing 1 to 5 carbons derived from ^13^C5-adenosine (designated as M+1 to M+5) between WT and KO cells (Figure 2L). These findings indicate that the increased whole-cell AMP levels in *Slc25a34*-KO BAC is likely derived from *de novo* purine synthesis.

To confirm that mitochondrial AMP predominantly originates from glycolysis, we isolated mitochondria from ^13^C6-glucose-labeled WT and *Slc25a34*-KO BAC and analyzed fluxes through the *de novo* purine synthesis pathway. Consistently, mitochondrial labeled AMP levels were higher, while cytosolic labeled AMP levels were lower, in *Slc25a34*-KO cells compared to WT. Additionally, ^13^C6-glucose-derived AMP was enriched in mitochondria (Figure 2M). Together, these experiments demonstrated that mitochondrial AMP is primarily derived from glucose by *de novo* synthesis. As such, ablation of *Slc25a34* results in the predominant accumulation of *de novo* synthesized AMP in the mitochondrial compartment.

One potential function of mitochondrially accumulated AMP could be to serve as a precursor for ADP and ATP synthesis. Our results showed that ADP and ATP also accumulated in mitochondria of *Slc25a34*-KO BAC, similar to AMP, raising the possibility that mitochondrial AMP might be converted to ADP and ATP (Figure S2G). However, labeled ADP and ATP derived from ^13^C6-glucose were unaltered or were even lower in both the mitochondrial and cytosolic compartments of *Slc25a34*-KO cells, indicating that the accumulated mitochondrial AMP is not used for ADP and ATP synthesis. Therefore, ADP and ATP accumulate in the mitochondria of *Slc25a34*-KO BAC may be attributed to the compensatory import of ADP for ATP production in response to the reduced whole-cell ATP levels (Figure S2H). Notably, mitochondrial DNA abundance in BAT remained unchanged in *Slc25a34*-KO mice (Figure S2I), suggesting that the mitochondrial AMP accumulation caused by *Slc25a34* deficiency may not be utilized for mitochondrial DNA synthesis either.

### SLC25A34 regulates BAT thermogenesis and mitochondrial stabilization in a cell-autonomous manner

Our results demonstrate that cold exposure induces Slc25a34 expression, enabling the transport of AMP from mitochondria to the cytosol in brown adipocytes. We next sought to investigate whether this cold-triggered AMP transportation is functionally linked to BAT thermogenesis. To explore the potential relationship between AMP levels and thermogenic function, we measured AMP abundance across different adipose tissue depots. Our analysis revealed that AMP is particularly enriched in BAT compared to two white adipose tissue depots. More importantly, BAT AMP levels were further upregulated in response to acute cold exposure (Figure 3A). In addition, after differentiation, AMP levels were higher in BAC than in 10T1/2 cells (Figure 3B). Furthermore, we revealed that *Slc25a34* expression is time-dependently upregulated by cold along with induced Ucp1 expression (Figure 3C). These results further support an association between AMP homeostasis and BAT thermogenic function.

**Figure 3.**
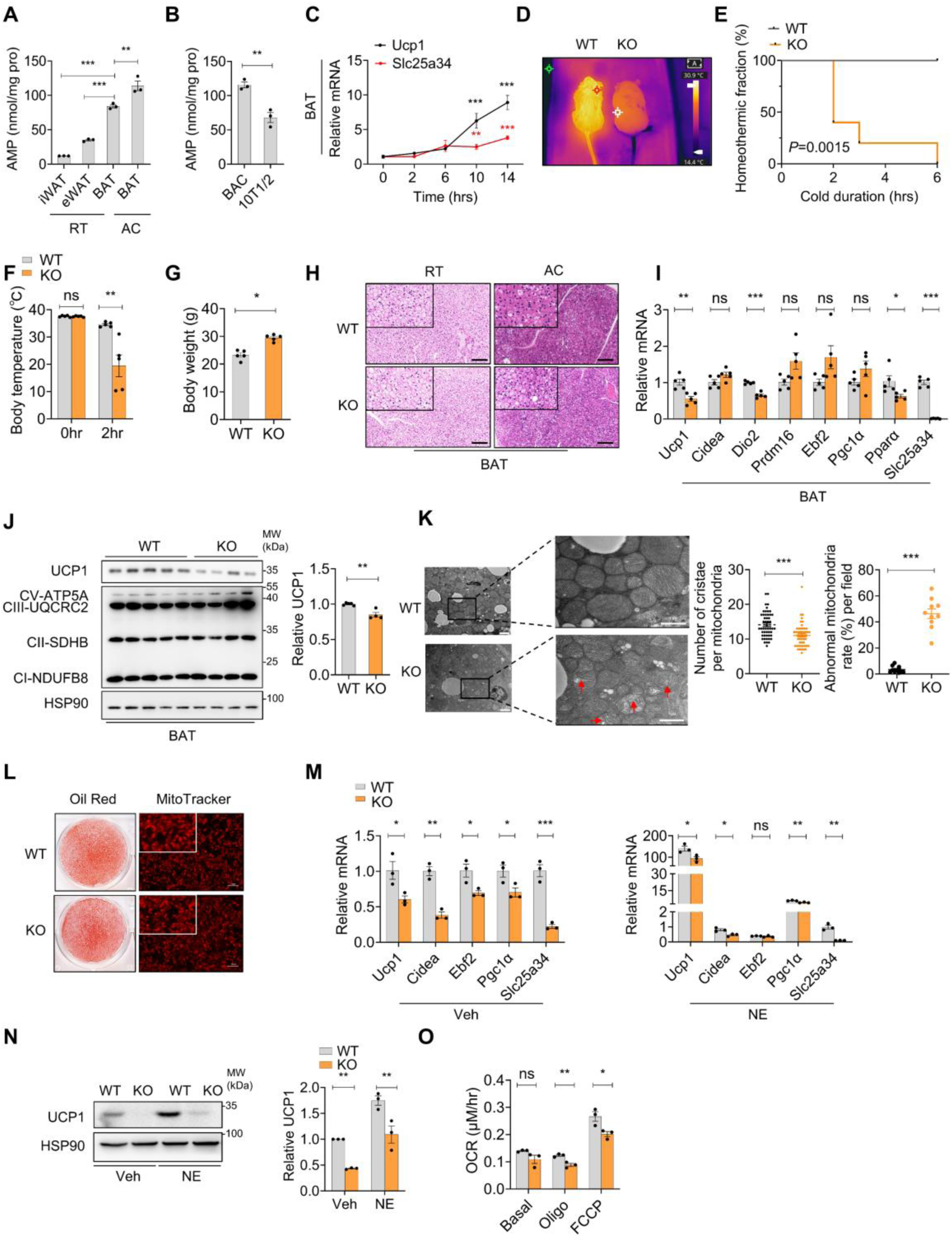
*Slc25a34*-KO mice exhibit impaired BAT thermogenesis and develops cold intolerance. (**A**) AMP concentrations were measured in BAT, eWAT, and iWAT from WT mice kept in room temperature, as well as in BAT from WT mice subjected to acute cold exposure at 4°C (n = 3). (**B**) AMP content in differentiated mouse brown adipocytes (BAC) and C3H10T1/2 (10T1/2) cells (n = 3). (**C**) QPCR analysis of mRNA expression of Slc25a34 and Ucp1 in brown adipose tissue (BAT) of mouse exposed to different duration of acute cold (4°C) (n = 4). (**D**) Infrared images of wild type (WT) and *Slc25a34*-knockout (KO) mice after exposed to acute cold (4°C, 6h). (**E**) The homeothermic fraction (rectal temperature>30°C) of Slc25a34-KO mice (n = 5) and WT mice (n = 5) in a 4°C acute cold exposure experiment. (**F**) Body temperature of *Slc25a34*-KO and control mice before and after 2 hours of cold exposure. (**G**) Body weight of Slc25a34-KO mice (n = 5) and WT mice (n = 5) after 4°C cold exposure. Hematoxylin and eosin (H&E) staining of brown adipose tissue (BAT) from WT and Slc25a34-KO mice housed at room temperature (RT) or subjected to acute cold exposure (AC) at 4°C. Scale bar=100 μM. (**I**) QPCR analysis of mRNA expression of thermogenic genes in the BAT of *Slc25a34*-KO (n = 5) and WT mice (n = 5) after acute cold exposure. (**J**) Protein from the BAT of *Slc25a34*-KO and WT mice used in **E** after cold exposure was extracted and analyzed via western blotting with the indicated antibodies (left). The relative amounts of UCP1 in left panel were determined by the density of the bands of UCP1, normalized to that of HSP90. Then, the fold changes were calculated by dividing the relative density of UCP1 in the Slc25a34-KO mice by that in WT mice (right). (**K**) Electron microscopy analysis of mitochondrial morphology in BAT from WT and *Slc25a34*-KO mice following 4°C cold exposure. Mitochondria exhibiting swelling and cristae disruption are highlighted with red arrows (left). Scale bar=1 μM. The number of cristae per mitochondria and the percentage of abnormal mitochondria was calculated (right) (**L**) WT and *Slc25a34*-KO brown adipocytes were isolated from mouse BAT and immortalized. Lipid droplets and mitochondrial content of WT and *Slc25a34*-KO BAC after differentiation were stained with Oil red O and Mitotracker, respectively. Scale bar=100 μM. (**M**) QPCR analysis of mRNA expression of thermogenic genes in differentiated WT and Slc25a34-KO BAC treated with vehicle (Veh) or norepinephrine (NE, 1 μM) for 6 hours (n = 3). (N) Western blot analysis of UCP1 protein in the differentiated WT and *Slc25a34*-KO BAC treated with vehicle (Veh) and NE (1 μM) for 6 hours (left). The relative amounts of UCP1 in left panel were determined by the density of the bands of UCP1, normalized to that of HSP90. Then, the fold changes were calculated by dividing the relative density of UCP1 in WT and Slc25a34-KO BAC treated with vehicle or NE by that WT BAC treated with vehicle (right). (**O**) Oxygen consumption rates of differentiated WT and *Slc25a34*-KO BAC (n = 3) were determined after FCCP (10 µM) and oligomycin (Oligo, 10 µg/ml) treatment. Data are presented as mean ± SEM. Statistical significance was determined by unpaired two-tailed Student’s t-test in (B), (F), (G), (I), (J), (K), (M), (N) and (O), one-way ANOVA multiple comparison test in (A), two-way ANOVA multiple comparison test in (C) and Log-rank (Mantel-Cox) test in (E). *P <0.05, **P <0.01, ***P <0.001. ns: not significant.

To assess the role of SLC25A34 in cold-induced thermogenesis, we performed an acute cold tolerance test in mice. Compared to WT controls, *Slc25a34*-KO mice displayed significantly heightened cold sensitivity (Figure 3D-E). Notably, KO mice developed severe hypothermia, with body temperatures dropping below 30°C within 2 hours of acute cold exposure (Figure 3F). The body weight of KO mice was greater after cold exposure (Figure 3G), whereas BAT weight was comparable between groups (Figure S3A). In addition, hematoxylin and eosin (H&E) staining revealed that lipid droplets in the BAT of WT mice were rapidly depleted following cold exposure, consistent with their utilization for thermogenesis, whereas lipid droplets persisted in *Slc25a34*-KO BAT, indicating impaired thermogenic function (Figure 3H). The expression of thermogenesis-related genes, including *Ucp1* and *Dio2*, was downregulated in the BAT of *Slc25a34*-KO mice compared to WT controls (Figure 3I). Consistent with the transcriptional changes, UCP1 protein levels were also reduced in *Slc25a34*-KO BAT. However, the expression of mitochondrial respiratory complex proteins remained unaltered (Figure 3J).

Acute cold exposure triggers mitochondrial biogenesis in BAT, a process that enhances respiratory capacity and supports thermoregulation. In our experiment, mitochondria in *Slc25a34*-KO mice exhibited swelling and abnormal fragmentation following cold exposure. Electron microscopy (EM) revealed reductions in cristae density per mitochondria and increase of abnormal mitochondria in *Slc25a34*-KO BAT compared to WT controls, further supporting the indispensable role of Slc25a34 in maintaining mitochondrial stability (Figure 3K).

To confirm that the impaired thermogenic phenotype in *Slc25a34* KO mice is attributable to *Slc25a34* deficiency in BAT, we rescued Slc25a34 expression in BAT via *in situ* injection of an *Slc25a34*-expressing adeno-associated virus (AAV) (Figure S3B). AAV-Slc25a34 administration restored *Slc25a34* expression in the BAT of KO mice, ablating the expression difference in BAT between WT and KO mice (Figure S3C). Following cold exposure, both WT and KO mice with BAT-specific Slc25a34 overexpression maintained normal body temperature, with no significant differences in body weight and BAT weight between groups (Figure S3D-F). Furthermore, lipid droplet accumulation in BAT was also unaltered (Figure S3G). Additionally, the expression of thermogenic genes was comparable between groups, as were the protein levels of UCP1 and mitochondrial respiratory complex subunits (Figure S3H-I). Collectively, these findings further establish a BAT-specific role for Slc25a34 in promoting adaptive thermogenesis in response to cold exposure.

To further investigate the role of SLC25A34 in brown and beige adipogenesis induced by the β3-adrenoreceptor agonist CL316,243 (CL), WT and *Slc25a34*-KO mice were intraperitoneally injected with CL for one week. Following treatment, Slc25a34 KO mice exhibited greater body weight, while no significant differences in BAT or iWAT weight were observed between groups (Figure S4A). H&E staining revealed that *Slc25a34*-KO BAT and iWAT had reduced multilocular lipid droplet structures, a hallmark of brown and beige adipocytes (Figure S4B). QPCR and western blot analyses confirmed that the absence of *Slc25a34* decreased the expression of thermogenesis-related genes at both the mRNA and protein levels in both BAT and iWAT following CL treatment (Figure S4C-F). Together, these findings indicate that Slc25a34 expression is required for β3-adrenergic signaling-mediated activation of brown and beige adipose tissues.

To further elucidate whether SLC25A34 regulates thermogenic gene expression in a cell-autonomous manner, we isolated and immortalized primary brown adipocytes from neonatal *Slc25a34*-KO mice and their WT littermate controls. Oil Red O and MitoTracker staining revealed no significant differences in adipogenesis and mitochondrial biogenesis in differentiated BAC between the two groups (Figure 3L). Notably, the expression of thermogenic genes, *Ucp1*, *Cidea* and *Pgc1α*, was significantly downregulated in *Slc25a34*-KO BAC under both vehicle and NE treatment conditions (Figure 3M), and UCP1 protein expression was consistently suppressed in *Slc25a34*-KO BAC (Figure 3N). Importantly, analysis of oxygen consumption rates (OCR) showed that following treatment with oligomycin and FCCP, OCR was significantly reduced in *Slc25a34*-KO BAC compared to WT controls (Figure 3O). These results demonstrate that Slc25a34 cell-autonomously regulates thermogenic gene expression and function in BAC.

### Slc25a34 mediates thermogenic gene expression by modulating AMPK signaling

AMP-activated protein kinase (AMPK), a central cellular energy sensor, monitors fluctuations in the AMP:ATP ratio. Upon activation, AMPK promotes BAT formation and increases thermogenic output ^17, 25, 26^. Cytosolic AMP levels are a key determinant of AMPK activation, thus we hypothesized that SLC25A34-mediated AMP transport might regulate cytosolic AMP levels and thereby influence AMPK activation.

Building on our previous finding of specific AMP enrichment in BAT, we further observed enhanced AMPK phosphorylation in BAT versus the other two white adipose depots (Figure 4A, Figure S5A). Notably, mature mouse BAC also exhibited more robust AMPK phosphorylation than 10T1/2 cells (Figure 4B, Figure S5B), and this trend was conserved in humans. Specifically, differentiated human BAC showed higher AMPK phosphorylation levels compared to their WAC counterparts (Figure 4C, Figure S5C). Additionally, acute cold exposure further increased AMPK phosphorylation in BAT, consistent with the elevated BAT AMP levels observed under cold conditions (Figure 4D, Figure S5D). To demonstrate the direct role of Slc25a34 in AMPK activation, we assessed AMPK phosphorylation in BAT from *Slc25a34*-KO and control mice following cold exposure and CL treatment. Loss of Slc25a34 significantly reduced phosphorylated AMPK levels, and this effect was recapitulated in *Slc25a34*-deficient brown adipocytes under both basal conditions and NE stimulation, indicating that Slc25a34 expression governs AMPK activation in brown adipocytes (Figure 4E–F, Figure S4D, S4F, S5E–F). Furthermore, re-expressing Slc25a34 in BAT normalized the difference in AMPK phosphorylation seen in *Slc25a34*-KO mice (Figure S3I), further validating that Slc25a34 regulates AMPK activation both in vitro and in vivo.

**Figure 4.**
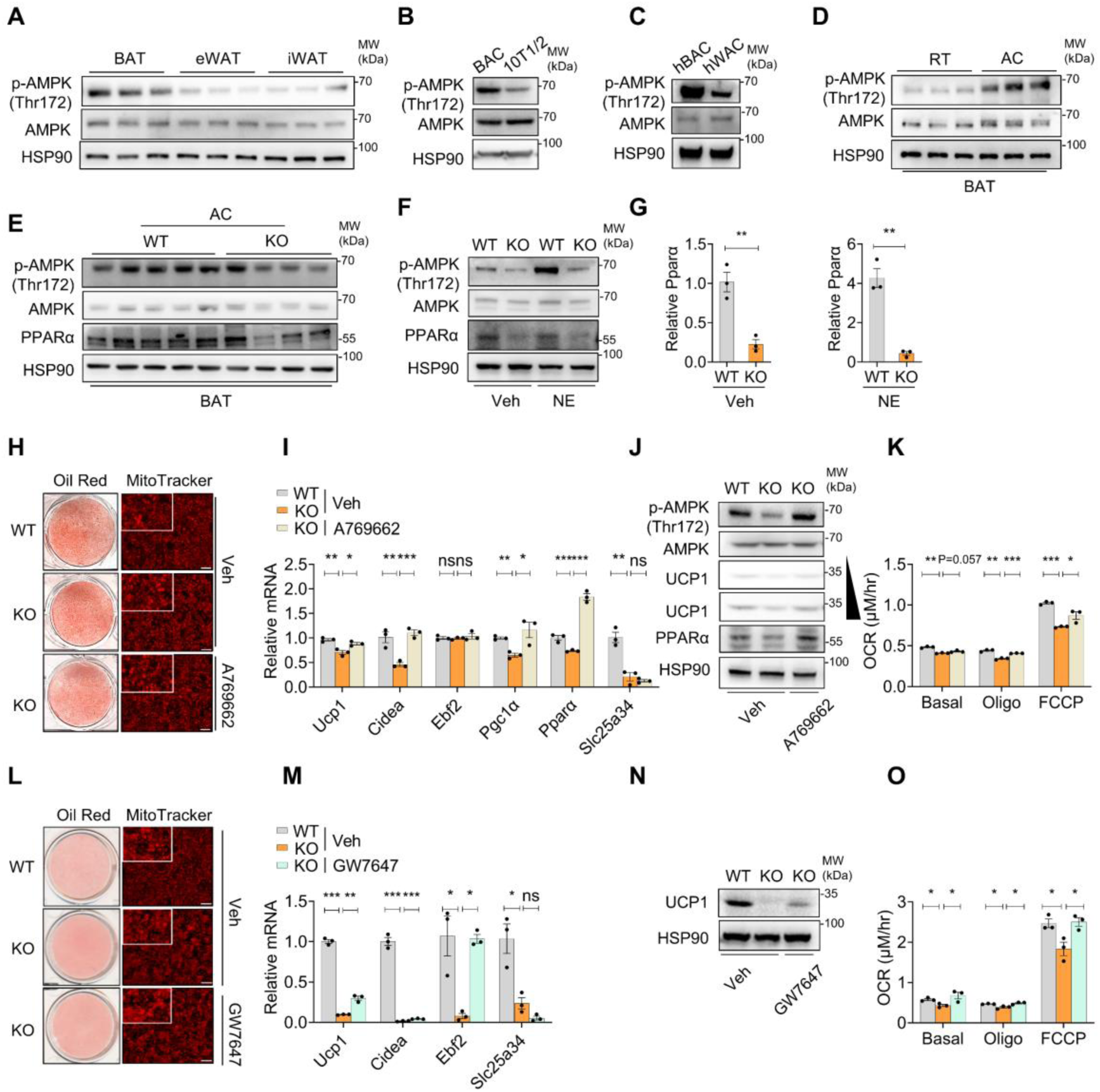
SLC25A34 mediates the AMPK cascade to enhance thermogenic gene expression and function. (**A**) Immunoblot analysis of phosphorylated and total AMPK in different fat depots from WT mice housed at room temperature (RT). (**B-C**) Immunoblot analysis of phosphorylated and total AMPK in differentiated mouse BAC and C3H10T1/2 (10T1/2) cells (B) and in differentiated human brown (hBAC) and white adipocytes (hWAC) (C). (**D**) Immunoblot analysis of phosphorylated and total AMPK in brown adipose tissue (BAT) from WT mice housed at RT or subjected to acute cold exposure at 4°C (AC). (**E**) Immunoblot analysis of indicated proteins in the BAT from WT and *Slc25a34*-KO mice subjected to acute cold exposure at 4°C (AC). (**F**) Immunoblot analysis of indicated proteins in the differentiated WT and Slc25a34-KO BAC (n = 3) treated with vehicle (Veh) or NE (1 μM, 6h). (**G**) QPCR analysis of Pparα expression in the differentiated WT and Slc25a34-KO BAC (n = 3) treated with vehicle (Veh) or NE (1 μM, 6h). (**H-K**) WT and *Slc25a34*-KO BAC were treated with vehicle (Veh) or A769662 (50 μM) during differentiation. (**H**) Lipid droplets and mitochondrial content were stained with Oil red O and Mitotracker respectively. Scale bar=100 μM. (**I**) Thermogenic gene expression was analyzed via QPCR (n = 3). (**J**) Protein was extracted and analyzed via western blotting with the indicated antibodies (**K**) Oxygen consumption rates (OCR) were measured after FCCP (10 µM) and oligomycin (Oligo, 10 µg/ml) treatment (n = 3). (**L-O**) WT and Slc25a34-KO BAC were treated with vehicle (Veh) or GW7647 (1 μM) during differentiation. (**L**) Lipid droplets and mitochondrial content were stained with Oil red O and Mitotracker respectively. Scale bar=100 μM. (**M**) Thermogenic gene expression was analyzed via QPCR (n=3). (**N**) Protein was extracted and analyzed via western blotting with the indicated antibodies. (**O**) Oxygen consumption rates (OCR) were measured after FCCP (10 µM) and oligomycin (Oligo, 10 µg/ml) treatment (n=3). Data are presented as mean ± SEM. Statistical significance was determined by unpaired two-tailed Student’s t-test in (G), and one-way ANOVA with dunnett’s test in (I), (K), (M) and (O). *P <0.05, **P <0.01, ***P <0.001. ns: not significant.

It has been reported that activated AMPK enhances the abundance and transcriptional activity of peroxisome proliferator-activated receptor alpha (PPARα), a key regulator of thermogenic gene expression ^27, 28^. Given our previous findings suggest that SLC25A34 mediates AMPK activation, we next investigated whether SLC25A34 regulates thermogenic gene expression through an AMPK-driven increase in PPARα signaling. Our results showed that *Pparα* mRNA and protein expression was significantly downregulated in the BAT of *Slc25a34*-KO mice following cold exposure (Figure 4E, Figure 3I). This effect was also observed in *Slc25a34*-KO BAC under both vehicle and NE treatment (Figure 4F-G). And restoring Slc25a34 in the BAT of *Slc25a34*-KO mice abolished the reduction in both Pparα mRNA and protein levels (Figure S3H-I).

To further confirm that AMPK-induced PPARα expression is required for SLC25A34-mediated induction of thermogenic gene expression and uncoupled respiratory function, we treated *Slc25a34*-KO BAC with the AMPK agonist A769662 and PPARα agonist GW7647 during differentiation. These experiments aimed to determine whether enhancing AMPK-PPARα signaling could rescue the impaired thermogenic gene expression and function in these *Slc25a34*-KO cells.

The results showed that A769662 treatment rescued the reduced AMPK phosphorylation in *Slc25a34*-KO cells, whereas this intervention exerted no effect on mitochondrial content or lipid droplet formation (Figure 4H, Figure S5G). Notably, in the presence of A769662, the expression levels of Pparα and thermogenic genes in *Slc25a34*-KO BAC were significantly restored (Figure 4I). A769662 treatment also rescued the protein expression of PPARα and UCP1 in *Slc25a34*-KO BAC (Figure 4J, Figure S5G). Furthermore, the defect in uncoupled respiration as indicated by OCR in *Slc25a34*-deficient BAC was also reversed by A769662 treatment (Figure 4K). Additionally, reactivating PPARα by GW7647 in *Slc25a34*-KO BAC also partially restored the expression of thermogenic genes and uncoupled respiration, with no impact on mitochondrial content or adipogenesis (Figure 4L-O, Figure S5H). These findings demonstrate that SLC25A34 activates the AMPK-PPARα cascade probably through AMP transport, thereby inducing thermogenic gene expression and function.

### SLC25A34 enhances glycolysis-TCA cycle flux to facilitate BAT thermogenesis

In contrast to WAT, BAT exhibits significantly higher rates of glucose uptake and utilization, particularly during thermogenesis ^29, 30^. Given that SLC25A34 has been proposed as a potential transporter of AMP, a known allosteric activator of phosphofructokinase (PFK), the rate-limiting enzyme in glycolysis, we hypothesized that SLC25A34 may enhance glycolysis and tricarboxylic acid (TCA) cycle flux to support BAT thermogenesis.

We first isolated cytosolic fraction, measured PFK expression and activity, and observed a significant reduction of PFK activity in *Slc25a34*-KO BAC (Figure 5A, Figure S5I). Additionally, analyses of glycolytic and tricarboxylic acid (TCA) cycle metabolites in WT and *Slc25a34*-KO BAC using metabolomic profiling revealed the levels of five glycolytic metabolites, including lactate, glucose 6-phosphate (G6P), pyruvate, thiamine pyrophosphate, and fructose 1,6-bisphosphate (F1,6BP), were reduced in *Slc25a34*-KO BAC compared to WT controls. Notably, this glycolytic impairment cascaded into the TCA cycle, leading to a decrease in four TCA cycle metabolites, aconitate, α-ketoglutarate, fumarate and malate (Figure 5B). Furthermore, ^13^C6-glucose tracing experiments revealed reduced incorporation of labeled glucose into downstream glycolytic and TCA cycle metabolites in *Slc25a34*-KO BAC (Figure 5C). These metabolites included M+3 lactate (containing 3 carbons from ^13^C6-glucose), M+2 succinate (containing 2 carbons from ^13^C6-glucose), and M+2 malate (containing 2 carbons from ^13^C6-glucose). Consistent with the specific enrichment of AMP in BAT, PFK expression and activity was also the highest in BAT relative to other white adipose depots in mice (Figure 5D, Figure S5J). Moreover, in *Slc25a34*-KO BAT following acute cold exposure, PFK activity was significantly reduced, concomitant with a decrease in lactate production (Figure 5E-F and S5K). These results suggest that SLC25A34 may regulate glycolytic-TCA cycle flux via PFK activation, thereby supporting thermogenesis during cold exposure.

**Figure 5.**
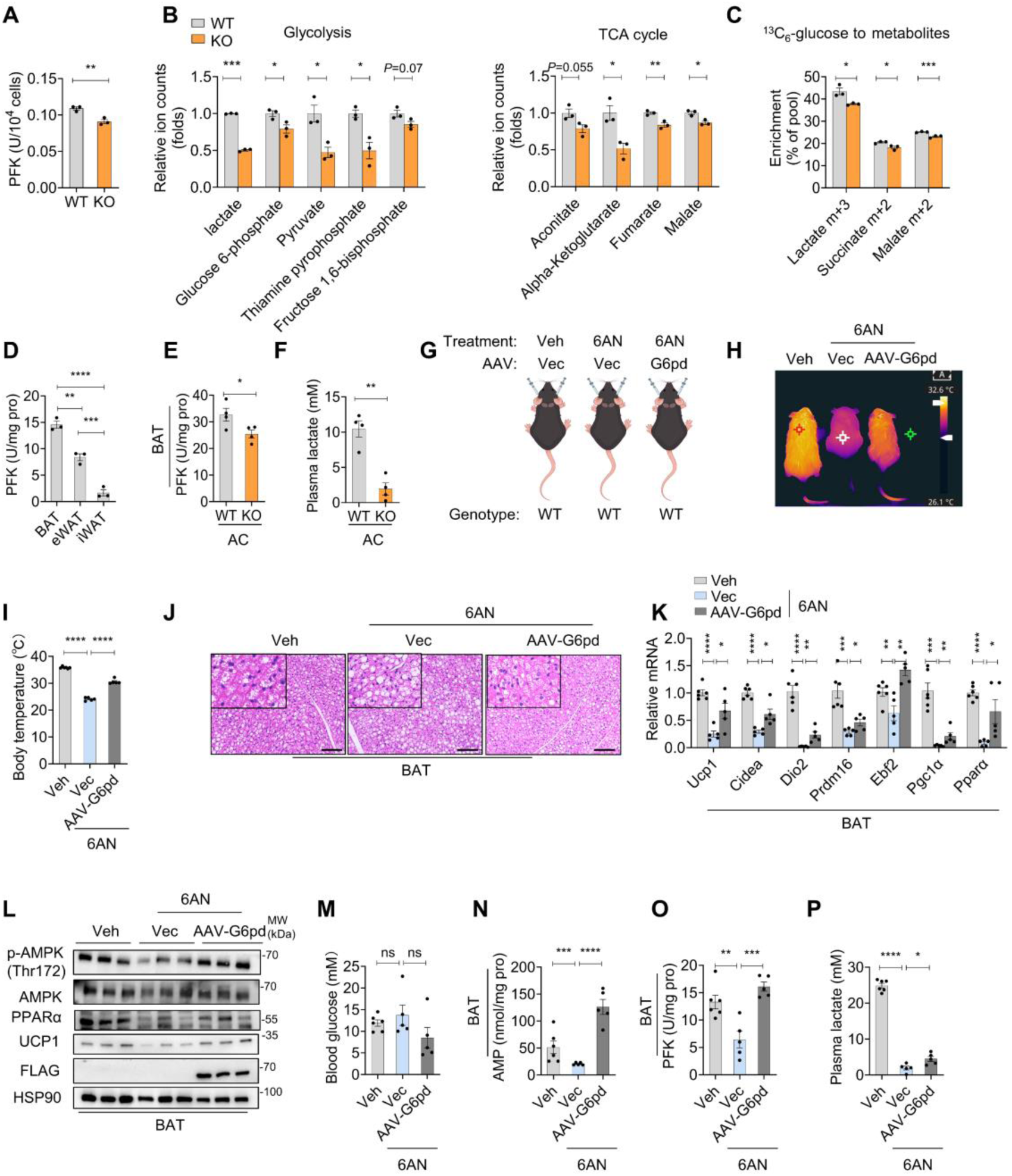
SLC25A34 regulates glycolysis-TCA cycle flux. (**A**) The activity of PFK was measured in differentiated wild type (WT) and *Slc25a34*-knockout (KO) mouse brown adipocytes (BAC, n = 3) and normalized to PFK protein expression. (**B**) Fold change of metabolites in glycolysis (left) and TCA cycle flux (right) in WT and Slc25a34-KO BAC (n = 3). (**C**) Mass isotopologue distribution analysis of lactate (m+3), succinate (m+2), and malate (m+2) in WT and Slc25a34-KO mouse BAC following 24-hour incubation with ^13^C6-glucose. (**D**) The activity of PFK was measured in different fat depots from WT mice housed at room temperature (RT, n = 3) and normalized to PFK protein expression. (**E**) The activity of PFK was measured in brown adipose tissue (BAT) of WT and Slc25a34-KO mice following 4°C cold exposure (AC, n = 4) and normalized to PFK protein expression. (**F**). The concentration of lactate was measured in plasma of WT and *Slc25a34*-KO mice following 4°C cold exposure (AC, n = 4). (**G**) Schematic of *in situ* BAT injection of vector (Vec) or G6pd AAV8 followed by 6-aminonicotinamide (6AN) treatment in WT mice housed at room temperature. (**H-P**) Mice were *in situ* BAT injected Vec or G6pd overexpression AAV, then treated with Vehicle or 6AN (10 mg/kg) for 5 consecutive days (Veh n = 6; 6AN n = 5; 6AN combined with G6pd AAV8 injection n = 5). Infrared thermal images (**H**) and body temperature (**I**) of mice. (**J**) Hematoxylin and eosin (H&E) staining of BAT of mice. Scale bar=100 μM. (**K**) Thermogenic gene expression was analyzed via QPCR. (**L**) Protein of BAT was extracted and analyzed via western blotting with the indicated antibodies. (**M**-**P**) The blood glucose (**M**), AMP content in BAT (**N**), PFK activity in BAT (**O**) and plasma lactate content (**P**) of mice. Data are presented as mean ± SEM. Statistical significance was determined by unpaired two-tailed Student’s t-test in (A), (B), (C), (E) and (F), and one-way ANOVA with dunnett’s test in (D), (I), (K), (M), (N), and (P). *P <0.05, **P <0.01, ***P <0.001. ns: not significant.

### De novo AMP synthesis is required for BAT thermogenesis

Our previous findings suggest that Slc25a34 mediates the transport of AMP from mitochondria to the cytosol, thereby promoting BAT thermogenesis during cold exposure. This raises the question of whether the *de novo* AMP synthesis pathway is required for BAT thermogenesis. To test this hypothesis, we administered 6-aminonicotinamide (6AN), an inhibitor of glucose-6-phosphate dehydrogenase (G6PD), the rate-limiting enzyme of the pentose phosphate pathway that supports AMP biogenesis, to a group of WT mice. In addition, we performed BAT-specific restoration of G6PD activity via *in situ* injection of a G6pd-expressing AAV into BAT, to verify the role of BAT-localized AMP synthesis in regulating BAT thermogenesis (Figure 5G). Our results showed that even when 6AN was administered at room temperature, the mice exhibited a robust hypothermic phenotype, with reduced body and BAT temperatures, alongside a slight reduction in body weight, while BAT weight remained comparable (Figure 5H-I, Figure S5L). Morphologically, the BAT of 6AN-treated mice has more lipid accumulation, and thermogenic gene expression in their BAT was also decreased (Figure 5J-L, Figure S5M). This impaired BAT thermogenesis phenotype was associated with reduced AMP levels, PFK activity, and lactate production whereas blood glucose levels remained unaltered (Figure 5M-P and S5N). More importantly, BAT-specific rescue of G6PD in 6AN-treated mice largely reversed the reductions in body and BAT temperatures, elevated BAT lipid content, attenuated thermogenic gene expression, and dysregulated AMP levels and PFK activity in BAT. Plasma lactate levels were also partially restored via AAV-mediated G6PD overexpression in BAT (Figure 5H-P, S5M-N). Collectively, these results indicate that *de novo* AMP synthesis in BAT is essential for its thermogenic capacity.

### PPARγ transcriptionally regulates Slc25a34 expression and subsequent AMP transport, thereby mediating AMPK activation and glycolytic activity

To identify the upstream regulator of *Slc25a34*, we performed a transcriptional regulator screening in BAC by overexpressing key thermogenic network transcription factors, including PPARγ, ZBTB7B, EBF2, C/EBPβ, PPARα, and PRDM16. Notably, PPARγ overexpression significantly upregulated *Slc25a34* expression in BAC (Figure S6A). To further validate this regulatory relationship, we treated BAC and 10T1/2 cells expressing vector control or PPARγ in combination with the PPARγ agonist rosiglitazone (Rosi). Rosi treatment increased *Slc25a34* expression. This induction is further augmented in the PPARγ overexpressing cells (Figure 6A). Conversely, treatment with T0070907 (T0907), a PPARγ antagonist, attenuated *Slc25a34* expression (Figure 6B). Moreover, *Pparγ* knockout in BAC markedly reduced *Slc25a34* expression (Figure 6C).

**Figure 6.**
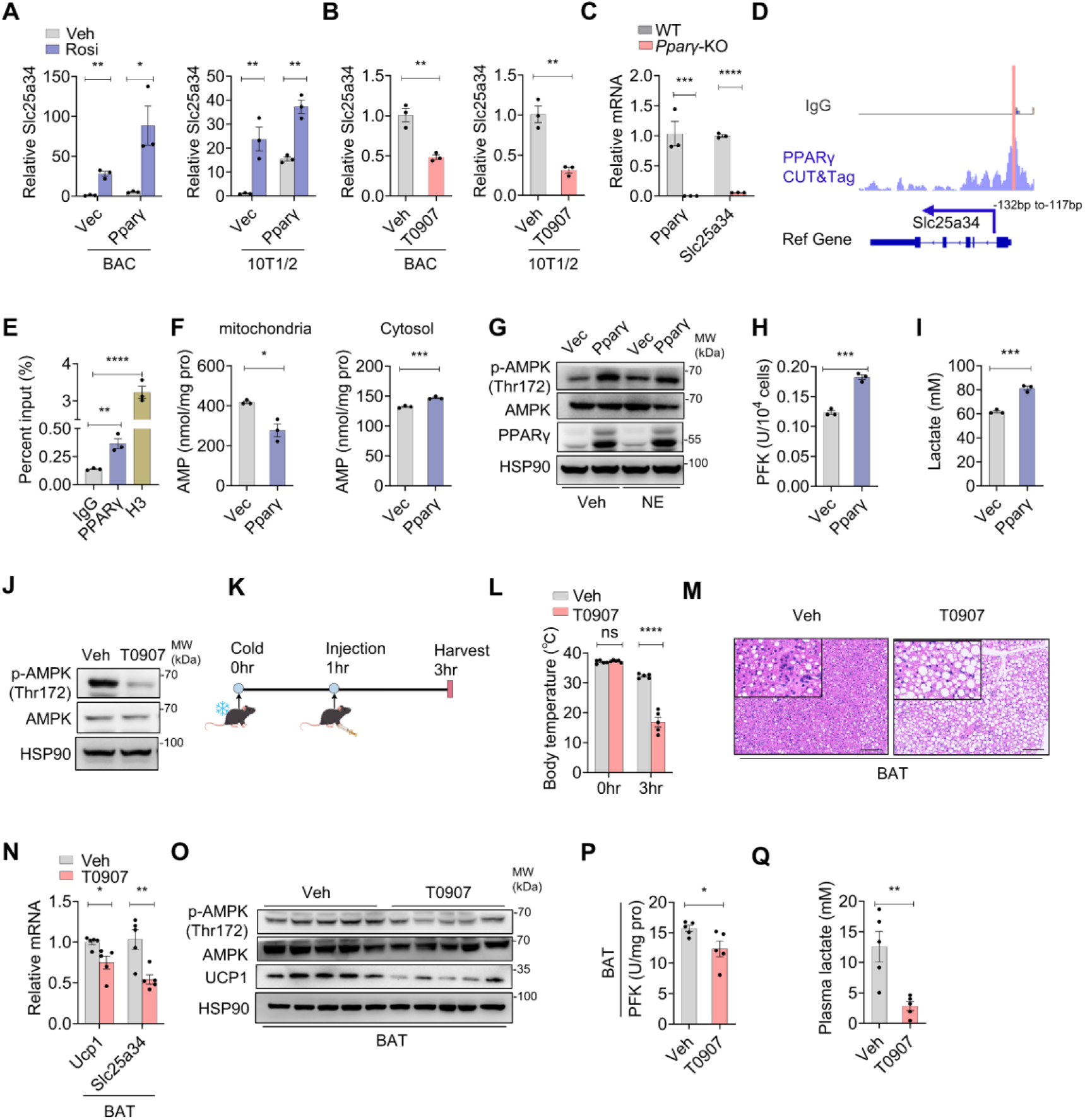
Slc25a34 is transcriptional induced by PPARγ. (**A**) QPCR analysis of Slc25a34 expression in differentiated mouse brown adipocytes (BAC) and C3H10T1/2 (10T1/2) cells overexpressing vector (Vec) or Pparγ (n = 3), treated with vehicle (Veh) or rosiglitazone (Rosi, 1 μM) during differentiation. (**B**) QPCR analysis of Slc25a34 expression in differentiated brown adipocytes (BAC) and C3H10T1/2 (10T1/2) cells following treatment with T0070907 (T0907, 10 μM) for 6 hours (n = 3). (**C**) QPCR analysis of Slc25a34 and Pparγ expression in differentiated WT (n = 3) and *Pparγ*-KO (n = 3) mouse BAC. (**D**) The binding activity of PPARγ to the promoter of Slc25a34 in differentiated mouse BAC was examined by Cut & Tag sequencing analysis. (**E**) ChIP-qPCR analyses were performed with anti-PPARγ antibody in differentiated PPARγ overexpressed mouse BAC. (**F**) AMP content in mitochondrial (left) and cytosolic fractions (right) of differentiated mouse BAC overexpressing PPARγ and vector (Vec) control (n = 3). (**G**) Immunoblot analysis of the indicated proteins in differentiated mouse BAC overexpressing PPARγ and vector (Vec) control, treated with vehicle or norepinephrine (NE, 1 μM) for 6 hours. (**H-I**) The PFK activity (**H**), normalized to PFK protein expression, and the lactate content (**I**) was measured in in differentiated mouse BAC overexpressing PPARγ and vector (Vec) control (n = 3). (**J**) Immunoblot analysis of phosphorylated and total AMPK in differentiated mouse BAC treated with Veh or T0907 (10 μM) for 6 hours. (**K**). Schematic diagram of the 4°C acute cold exposure experiment with T0907 treatment. (**L**) Body temperature of Vehicle- (Veh-) (n = 5) and T0907- (50mg/kg) (n = 5) treated mice before and after 2 hours of cold exposure. (**M**) Hematoxylin and eosin (H&E) staining of BAT from mice used in **L**. Scale bar=100 μM. (**N**) QPCR analysis of Slc25a34 and Ucp1 expression in BAT of mice used in **L**. (**O**) Immunoblot analysis of protein indicated in BAT of mice used in **L**. (**P-Q**) The activity of PFK in BAT (**P**), normalized to PFK protein expression and the plasma lactate content (**Q**) of mice used in **L**. Data are presented as mean ± SEM. Statistical significance was determined by unpaired two-tailed Student’s t-test in (A), (B), (C), (F), (H), (I), (L), (N), (P), and (Q), and one-way ANOVA with dunnett’s test in (E). *P <0.05, **P <0.01, ***P <0.001. ns: not significant.

To elucidate whether PPARγ directly binds to the *Slc25a34* promoter, we constructed a *Slc25a34* promoter luciferase reporter containing a mutated PPAR response element (PPRE) binding site. Luciferase activity assays were performed in HEK293T cells co-transfected with the *Slc25a34* promoter reporter and a PPARγ overexpression plasmid (Figure S6B). The results revealed that the PPAR response element (PPRE) binding site, located in the region spanning -132 bp to -117 bp relative to the transcriptional start site (TSS), is critical for the PPARγ-mediated transcriptional regulation of *Slc25a34* (Figure S6C). Furthermore, CUT&Tag-seq and PPARγ Chromatin Immunoprecipitation (ChIP) assays confirmed the direct binding of PPARγ to the *Slc25a34* promoter within the -132 bp to -117 bp region (Figure 6D-E). Together, these findings demonstrate that PPARγ binds to the “-132 bp to -117 bp” PPRE in the *Slc25a34* promoter to drive its expression.

To further investigate whether PPARγ regulates *Slc25a34* expression and thereby mediates AMP transport between mitochondria and the cytosol, we measured mitochondrial and cytosolic AMP levels in BAC overexpressing PPARγ. Our results demonstrated that PPARγ overexpression enhanced AMP efflux from mitochondria (Figure 6F). Interestingly, PPARγ overexpression in BAC also stimulated the phosphorylation of AMPK (Figure 6G, Figure S6D). Additionally, PPARγ overexpression increased PFK activity and lactate production in BAC, indicating that PPARγ promotes glycolytic flux (Figure 6H-I, Figure S6E). In contrast, inhibition of PPARγ with T0907 suppressed AMPK phosphorylation (Figure 6J, Figure S6F). Thus, these findings indicate that PPARγ expression modulates SLC25A34-mediated AMP efflux to the cytosol, thereby triggering AMPK activation and boosting glycolytic activity.

To further investigate whether PPARγ induces *Slc25a34* expression during cold stimulation, we administered the PPARγ antagonist T0907 to mice during acute cold exposure (Figure 6K). Following 3 hours of cold exposure, T0907-treated mice developed a hypothermic phenotype with lowered body temperature (Figure 6L). This was associated with morphological whitening of BAT and a pronounced decrease in Slc25a34 expression in BAT, whereas these mice exhibited modestly higher body weight and comparable BAT weight relative to controls (Figure 6M-N, Figure S6G). Notably, T0907 treatment impaired the cold-induced phosphorylation of AMPK in BAT and reduced UCP1 expression (Figure 6N-O, Figure S6H), and inhibited both PFK activity (Figure 6P, Figure S6I) of BAT and lactate production (Figure 6Q). These findings support that PPARγ mediates the upregulation of *Slc25a34* during cold exposure and this process is essential for cold-induced BAT thermogenesis.

## Discussion

The SLC25A family of mitochondrial transporters plays essential roles in mediating metabolic crosstalk between mitochondria and the cytosol. Individual members of this family govern the transport of nucleotides, amino acids, and organic acids, thereby regulating cellular metabolism and influencing disease states ^32, 33^. However, whether these transporters can sense environmental stimuli and contribute to specific physiological processes remains unclear. Cold-induced thermogenesis is a critical physiological process for maintaining body temperature and energy homeostasis in mammals. Nevertheless, the underlying mechanisms by which cells perceive cold stimuli and convert them into metabolic adaptations remain incompletely understood. In this study, we reveal a novel signaling pathway that links mitochondrial-cytosolic metabolite transport to BAT activation. Specifically, we identify SLC25A34, a member of the mitochondrial solute carrier family, as a cold-responsive transporter that mediates the efflux of AMP from mitochondria to the cytosol. This transport event acts as a key metabolic signal, activating AMPK signaling and glycolysis to ultimately promote thermogenesis.

Through systematic screening of SLC25A family genes in BAT, we identified Slc25a34 as a robustly cold-induced transporter, with its expression tightly correlated with thermogenic activation. Genetic ablation of *Slc25a34* impaired cold tolerance, compromised mitochondrial stabilization and reduced BAT thermogenesis, establishing SLC25A34 as a key regulator of adaptive thermogenesis.

Mechanistic studies revealed that SLC25A34 functions as a dedicated mitochondrial AMP exporter. Several lines of evidence support this conclusion. First, metabolomic profiling of *Slc25a34*-deficient BAC revealed significant mitochondrial AMP accumulation concurrent with cytosolic AMP diminishment, a phenotype indicative of impaired AMP export. This was further confirmed by visualization of AMP localizing to mitochondria following *Slc25a34* depletion. Second, direct binding assays demonstrated specific interaction between SLC25A34 and AMP, with no detectable affinity for other adenine nucleotides. Third, the structural prediction by AlphaFold identified a high-quality binding pocket for AMP in SLC25A34, consistent with its substrate specificity. More importantly, the proteoliposome assay further confirmed that SLC25A34 is capable of AMP transport. These findings identify SLC25A34 as the first characterized mitochondrial AMP transporter, expanding the functional repertoire of the SLC25A family beyond its known roles in metabolites transport.

Cytosolic AMP is a well-characterized metabolic signal that activates AMPK and enhances glycolysis via allosteric regulation of PFK, thereby coordinating energy stress responses and metabolic flux. Recent study shows that AMP could also bind to NAMPT to mediate NAD production ^34^. In contrast, the role of mitochondrial AMP has been largely overlooked. Our study reveals that mitochondrial AMP serves as a specialized signaling reservoir. Compared to metabolites like citrate, which shuttle bidirectionally, we revealed that under basal conditions, mitochondrial AMP levels are substantially higher than those in the cytosol, with cold exposure triggering its rapid release into the cytosol via SLC25A34. Notably, our metabolite tracing experiments exclude major roles for mitochondrial AMP in canonical mitochondrial processes that converted AMP to ADP/ATP for energy production nor utilized for mitochondrial DNA synthesis via dNTP generation. Instead, we propose that mitochondrial AMP functions primarily as a stored signaling molecule, enabling rapid activation of cytosolic pathways in response to environmental stimuli such as cold exposure. This storage mechanism likely confers a significant kinetic advantage. Specifically, it enables BAT to mount a rapid thermogenic response without relying on *de novo* transcriptional cascade induction during acute cold exposure.

We further demonstrate that mitochondrial AMP is predominantly derived from glucose via *de novo* purine synthesis. This links nutrient availability to thermogenic capacity, with *Slc25a34* deficiency triggering compensatory upregulation of *de novo* AMP synthesis, likely to restore cytosolic AMP levels. More importantly, inhibiting *de novo* AMP synthesis through the blockade of G6PD activity potently attenuates BAT thermogenesis. These findings highlight *de novo* purine synthesis as a rate-limiting step in cold-induced BAT activation, positioning mitochondrial AMP synthesis and export as integral components of metabolic reprogramming in thermogenesis.

AMPK is a master regulator of energy homeostasis, activated under low energy conditions to promote catabolic pathways and thermogenic gene expression via induction of PPARα and downstream targets like UCP1 ^35, 36^. Our data establish SLC25A34 as a novel upstream activator of AMPK in cold-exposed BAT. This conclusion is supported by two key observations: first, Slc25a34 expression and AMPK phosphorylation increase in parallel during cold exposure, and second, *Slc25a34* knockout significantly abolishes cold-induced AMPK activation. Importantly, pharmacological activation of AMPK rescues uncoupled respiration defects in *Slc25a34*-deficient BAC, confirming AMPK as a critical downstream effector of SLC25A34-mediated signaling. This places SLC25A34 as a critical link between mitochondrial metabolism and AMPK-dependent thermogenesis.

Beyond AMPK, SLC25A34-mediated AMP export enhances glycolytic flux via allosteric activation of PFK, a key regulatory enzyme in glycolysis. This dual regulation, activating both AMPK signaling and glycolysis, creates a synergistic mechanism to boost energy expenditure: AMPK drives thermogenic gene expression, while increased glycolysis provides substrates for mitochondrial respiration and heat production. Together, these data reveal a novel mechanism that metabolite transport coordinates multiple metabolic and signaling pathways to support thermogenesis.

Transcriptional networks, particularly those governed by PPARγ, are central to BAT development and thermogenic function. Our data reveal that cold exposure promotes PPARγ binding to the *Slc25a34* promoter, driving its expression. This establishes a critical axis wherein cold-induced PPARγ activates Slc25a34, thereby enhancing mitochondrial AMP export to trigger AMPK activation, which in turn amplifies the expression of thermogenic genes. Disruption of this axis via *Slc25a34* knockout or PPARγ inhibition impairs cold tolerance, underscoring the functional importance of this PPARγ-SLC25A34-AMPK axis in adaptive thermogenesis. This axis integrates transcriptional regulation with metabolic signaling, ensuring that cold-induced transcriptional changes are tightly coupled to metabolic outputs. Such coordination likely optimizes thermogenic efficiency, allowing BAT to scale its activity in response to environmental cold.

In summary, our study identifies a novel cold-responsive metabolic signaling pathway in which SLC25A34-mediated mitochondrial-to-cytosolic AMP transport activates AMPK and glycolysis to drive BAT thermogenesis. This pathway highlights mitochondrial AMP as a specialized signaling reservoir, links *de novo* purine synthesis to thermogenic capacity, and identifies a PPARγ-SLC25A34-AMPK regulatory loop that coordinates transcriptional and metabolic adaptations. These findings advance our understanding of how subcellular metabolite transport governs cellular function and underscore the potential of targeting mitochondrial metabolite transporters to enhance thermogenesis in metabolic disease.

### Limitations and future directions

While our study identifies SLC25A34 as a key regulator of cold-induced thermogenesis, several questions remain. First, the mechanism by which AMP is transported into mitochondria for storage remains unknown; identifying the mitochondrial AMP importer will be critical to understanding the complete AMP trafficking cycle. It is also possible that mitochondrial AMP is generated from ADP via mitochondrially localized adenylate kinase. Second, the role of SLC25A34 in other metabolically active tissues for example skeletal muscle and liver, and its potential involvement in other physiological contexts such as exercise and fasting warrant exploration. Finally, preclinical studies with SLC25A34 modulators will be necessary to validate its therapeutic potential in obesity.

## Methods

### Animal experiments

Our study examined male mice in chow-diet -feeding and acute cold exposure experiments because male animals exhibited less variability in phenotype. All animal studies were conducted following the guidelines outlined in the Animal Care and Use Committee (IACUC) of Shanghai Jiao Tong University School of Medicine (No. A2022-064). Mice aged between 8 and 28 weeks were used in the experiments. *Slc25a34*-KO mice were bought from GemPharmatech Co. Ltd in a C57BL/6J background. All mice were provided with regular rodent chow and maintained under a 12-hour light/dark cycle. To activate β3-AR, CL316,243 (1 mg/kg, C5976, Sigma-Aldrich) was administered intraperitoneally daily for 7 days. To inhibit *de novo* AMP synthesis pathway, 6-aminonicotinamide (6AN,10 mg/kg, HY-W010342, MCE) was administered intraperitoneally daily for 5 days. For acute cold exposure experiment, mice were maintained at room temperature (∼25°C) before cold exposure. During cold exposure, the mice were maintained in a temperature-controlled chamber at 4°C with free access to food and water. Rectal temperature was measured with an electronic thermometer (ALC-CT, ALCBIO) every 2 hours during cold exposure. Thermal infrared images were captured using an infrared camera (M300, InfiRay). We set a rectal temperature lower than 30°C as the cutoff to calculate the homeothermic fraction. For cold acclimation, mice were placed in a temperature-controlled chamber at 22 °C and subjected to a gradual decrease in ambient temperature (4 °C per day) to a final temperature of 10 °C, which was maintained for more 5 days before tissues were harvested. The PPARγ antagonist T0070907 (HY-13202, MCE) was administered intraperitoneally at a dose of 50 mg/kg, 1 hour after acute cold exposure initiation, during cold challenge. After acute cold exposure, mice were euthanized and tissues were dissected, promptly frozen in liquid nitrogen before the extraction of RNA and protein. Plasma was stored at -80°C after centrifugation until further processing. For histological analysis, tissues were dissected, fixed in 4% formalin overnight at 4°C, and then embedded in paraffin followed by H&E staining. All experiments were independently repeated and performed at least twice. Samples or data points were excluded solely due to technical equipment or human errors that led to poor control of a sample.

### Cell culture and differentiation

HEK293T cells and C3H/10T1/2 cells were obtained from American Type Culture Collection (ATCC). Mouse brown preadipocytes were isolated from the interscapular brown adipose tissue (BAT) of newborn wild-type (WT) or *Slc25a34*-knockout (KO) mice; The cells were immortalized by expressing the large T antigen of the SV40 virus as previously reported ^37^. Briefly, the coding sequence of SV40 large T antigen was cloned into the retroviral transfer plasmid (pMSCV-plus-hygromycin) and co-transfected with packaging plasmids (Gag-pol and VSVG) into 293T cells to acquire the pseudoretrovirus. Primary mouse brown preadipocytes were infected with these retroviruses to introduce the large T antigen, and hygromycin selection was performed to obtain stable immortalized cells. Human primary brown and white preadipocytes were isolated from fat tissue obtained from the area near the supraclavicular region in the neck^38^ or from the subcutaneous region, which was obtained from Shanghai Ninth People’s Hospital. All human tissue collection protocols were approved by the Ethics Committee of Shanghai Ninth People’s Hospital. HEK293T cells, C3H/10T1/2 cells, mouse brown preadipocytes and human brown and white preadipocytes were maintained in DMEM supplemented with 10% fetal bovine serum (Sigma, F8318). Confluent C3H/10T1/2 cells, mouse brown preadipocytes and human preadipocytes were subjected to differentiation by adding induction medium containing 0.5 mM IBMX (ACROS, 228420050), 125 μM indomethacin (Sigma, I-7378), 1 μM dexamethasone (Sigma, D2915), 20 nM insulin (Sigma, I5500) and 1 nM T3 (Sigma, T2877). Cells were switched to differentiation medium (DMEM, 10% FBS, 20 nM insulin and 1 nM T3) after two days. The differentiation medium was changed every two days until mature adipocytes formed. The cells were treated with vehicle or 1 μM norepinephrine (Sigma, A9512) for 6 hours to activate the expression of thermogenic genes before being collected. To manipulate PPARγ and AMPK activity, we incubated BAC with PPARγ agonist, Rosiglitazone (Rosi, 1 μM, HY-17386, MCE), and antagonist, T0070907 (T0907, 10 μM, HY-13202, MCE) and the AMPK agonist A769662 (50 μM, HY-50662, MCE), and the PPARα agonist GW7647 (1 μM, HY-13861, MCE) throughout the differentiation process.

### Mitotracker and Oil-Red-O staining

The mitochondrial contents of the cells were stained with MitoTracker probe (Invitrogen, MP07510) according to the manufacture’s instruction. Briefly, the cells were washed once with serum-free DMEM and incubated with MitoTracker probes (250 nM) in serum-free DMEM incubated for 30 minutes at 37°C. Then, the cells were washed with serum-free DMEM three times, and fluorescence images were obtained via fluorescence microscope. For Oil Red O staining, after brown adipocyte differentiation, the cells were fixed with 2% formalin for approximately 30 minutes at room temperature. Then, the cells were rinsed three times with distilled water and stained with a 60% Oil Red O (O0625, Sigma) solution for 2 hours at room temperature. Next, the cells were washed three times with double distilled water and then photographed after air dry.

### Raman imaging

Raman imaging in differentiated WT and *Slc25a34*-KO BAC sections was conducted with a confocal Raman microscope WITec Alpha300R (WITec GmbH, Ulm, Germany) equipped with a 532 nm laser set to the power of 30 mW in front of the 50x objective (NA0.75). Scans were acquired with 0.08 s integration time in areas sized 22 × 22 μm with a resolution of 350 nm (180000 spectra per scan). Laser power and integration time were optimized to avoid burning of the tissue. Similar parameters were also employed by other groups. Nevertheless, each scan was checked for thermal damage artifacts, observable by changes in peak position and width, after imaging in the WITec Project 7.1 software (WITec GmbH, Ulm, Germany) to ensure data quality. Colocalization of AMP and CytC were quantified by using ImageJ (v.1.53) and Colocalization Finder (v.1).

### Immunoblotting analysis

RIPA buffer (150 mM NaCl, 1% NP40, 50 mM Tris-HCl (pH 8.0), 25 mM NaF, 2 mM Na3VO4, 1 mM PMSF, and protease inhibitor cocktail) was used to lyse cells or tissues, The concentration of protein was determined by using detergent compatible Bradford protein assay kit (P0006C, Beyotime). Lysate containing 30 μg of whole protein were mixed with a 2 x SDS-PAGE reduced loading buffer and heated at 98°C for 5 minutes for each sample. The procedure for western blotting was followed as described previously ^41^. Briefly, proteins were separated via SDS-PAGE and transferred onto PVDF membranes. The membranes were blocked with 5% non-fat milk in TBST (Tris-buffered saline with 0.1% Tween-20) for 1 hour at room temperature, followed by incubation with primary antibodies against UCP1 (UCP11-A, Alpha Diagnostic), SLC25A34 (17557-1-AP, Proteintech), AMPKα (#2532, CST), p-AMPKα (Thr172) (#2535, CST), PPARα (ab126285, Abcam), HSP90 (1317-1-AP, Proteintech), OXPHOS (ab110413, Abcam), PPARγ (#2443, CST), PFK (68385-1-Ig, Proteintech) and FLAG (M185-3L, MBL) overnight at 4°C. After TBST washing 5 times while agitating for 5 minutes per wash, membranes were incubated with HRP-conjugated secondary antibodies (115-035-003, 111-035-003, Jackson) for 1 hour at room temperature and washed 5 times in TBST followed by signal detection using an ECL kit (MA0186-2, Mellunbio).

### Gene expression analyses

Total RNA was extracted from tissue specimens or cultured cells using TRIzol reagent (R401, Vazyme) in accordance with the manufacturer’s instructions. RNA purity and concentration were assessed using MULTISKAN Sky microplate reader (Thermo Fisher Scientific). For cDNA synthesis, 1 µg of total RNA underwent reverse transcription with the HiScript II Q RT SuperMix (R222-01, Vazyme). Quantitative real-time PCR (qPCR) was executed on a Roche LightCycler 480 II platform using SYBR Green (Q311, Vazyme). Relative gene expression levels were normalized to the housekeeping gene AP0. Primers for qPCR were listed in Table S1.

### Adeno-Associated Virus (AAV) construction and *in situ* BAT administration

AAV-CAG-MCS plasmid served as target vectors for constructing AAV. The coding sequence of Slc25a34 and G6pd was amplified from cDNA of BAT from wild type C57BL/6 mouse and insert into the AAV-CAG-MCS plasmid (AAV-Slc25a34 and AAV-G6pd1). Green fluorescent protein (GFP) was inserted into AAV-CAG-MCS act as vector control (AAV-GFP). For AAV production, the recombinant AAV plasmid was co-transfected with the pAdΔF6 and pAAV8 serotype plasmids into HEK293T cells using polyethylenimine (PEI, MX2202, Shanghai Maokang). After 60 hours, cells were harvested and the AAV particles were purified using iodixanol gradient ultracentrifugation in OptiSeal tube at 53,000 rpm and 14°C for 3 hours. The viral titer was determined by quantitative PCR (qPCR). To achieve Slc25a34 and G6pd overexpression in the BAT, AAV was bilaterally administered into the interscapular BAT depots. Mice were anesthetized with isoflurane, and the fur over the scapular region was removed. A horizontal skin incision was made to expose the underlying BAT pads, which were gently exteriorized using forceps. Subsequently, 50 μl of viral suspension containing 5 × 10^11^ viral genomes (vg) was injected into each fat pad at 5–8 distinct sites to ensure uniform distribution. The incision was then closed using surgical staples. The mice injected with AAV were received subsequent treatment after 7 days.

### Hematoxylin-eosin (H&E) staining

Following euthanasia, mice tissue such as BAT, iWAT, eWAT were dissected and fixed in 4% paraformaldehyde and processed through a dehydration series using ethanol gradients. The tissues were then embedded in paraffin and sliced into 4 μm sections. Dimethylbenzene was used to dewax the slides, which were then rehydrated through a gradient of ethanol concentrations. H&E staining was carried out according to the instructions of the Hematoxylin-eosin constant dye kit. Following staining, the slides were dehydrated with absolute ethanol, normal butanol, and dimethylbenzene, and subsequently mounted using Rhamsan gum. The stained tissues were examined microscopically.

### Separation of Stromal and vascular fraction (SVF) and adipocyte fraction

Five mice were used for SVF and adipocyte fraction isolation. Briefly, interscapular brown adipose tissue was excised and kept in ice-cold HBSS (B430KJ, BasalMedia) until all BAT was collected. Then BAT was transferred to a new cell culture dish, chopped into small pieces using scissors for 2 minutes and resuspended with 2.5 mL 2 x digestion buffer (HBSS supplemented with 1% BSA (36104ES25, Yeasen), 6.25 mM CaCl₂, 5 mM glucose, and 1 μM adenosine (164040050, Acros), and transferred into a 50 mL conical tube. The mixture was supplemented with an additional 2.5 mL of 2 x collagenase solution (Serum free high-glucose DMEM containing 1% BSA, 2.4 U/mL dispase II (4942078001, Roche), 3 mg/mL Type I collagenase (LS004196, Worthington), and 10 mM CaCl₂), mixed well and incubated for 30 minutes at 37 °C while shaking (150 rpm/min) at water bath. During incubation, tubes were inverted and gently shaked every 5 minutes. Following the complete digestion, the whole digestive mixture was filtered through 100 µm cell strainer (Biofil) into 15ml conical tube. 5 ml DMEM containing 10% FBS was employed to stop the digestion and rinse the filter. The tubes of mixture were centrifuged at 1000rpm for 5 minutes to pellet SVF. The floating fraction was delicately moved into a new tube using a wide opening 1 ml pipet. The fat fractions were acquired after the liquid was sucked away by the syringe. For SVF pellet, the supernatant was discard and washed with DMEM once and spin at 1000 rpm/min for 5 minutes to pellet SVF again. The SVFs and adipocyte fractions were used for RNA extraction or ATP/ADP/AMP detection immediately.

### Transmission electron microscopy

Mice were euthanized following 6 hours of acute cold exposure, and BAT was dissected and cut into 1-2 mm pieces. The tissue was fixed overnight at 4°C in a solution containing 2.5% glutaraldehyde and 2% paraformaldehyde, then post-fixed in 1% osmium tetroxide. Samples were dehydrated through a graded ethanol series (30%, 50%, 70%, 80%, 95%, and 100%), with a 10-minute incubation at each concentration. Next, samples were infiltrated with Embed 812 resin for 6 hours, transferred to capsules filled with Embed 812, and polymerized at 60°C for 48 hours. Ultrathin sections (70-90 nm thick) were cut, stained with uranyl acetate and lead citrate to enhance contrast, and prepared for imaging. Observations were made using a transmission electron microscope. Abnormal mitochondria were counted and the morphometric evaluation of mitochondria and cristae were outlined using Image J software.

### Luciferase reporter assay

The mouse *Slc25a34* promoter sequence (-1875 bp to +136 bp) was cloned into the pGL3-Basic vector (Promega) to construct a *Slc25a34* promoter-containing luciferase plasmid (*Slc25a34*-luc). Predicted PPARγ binding sites, located at positions -132 bp to -117 bp and -682 bp to -667 bp relative to the TSS, were mutated using the Mut Express II Fast Mutagenesis Kit (Cat. No. V2-C214, Vazyme) in accordance with the manufacturer’s instructions. These binding sites were initially identified via the PPARgene database (http://www.ppargene.org/). Luciferase reporter assays were performed in HEK293 cells, as previously described ^42^. Briefly, cells were co-transfected with 50 ng of the *Slc25a34* -luc plasmid plus either empty vector or 100 ng of a PPARγ expression construct. Relative luciferase activity was measured 48 hours post-transfection using luciferase substrate (Promega, E1500) and measured via a microplate reader system (Molecular Devices, ID5), all experiments were conducted in triplicate.

### ChIP-qPCR

Chromatin immunoprecipitation was performed in differentiated BAC cells using the SimpleChIP® Enzymatic Chromatin IP Kit (Magnetic Beads) (#9003,CST) following the instructions. To cross-linking proteins to DNA, differentiated BAC cells were incubated with culture medium supplemented with 1% formaldehyde at 37°C for 10 minutes. The reaction was quenched using 0.125 M glycine for 5 minutes at room temperature and subsequently washed with ice-cold PBS containing protease inhibitor cocktails (PIC). 2ml ice-cold PBS containing PIC was added. Cells was then scraped and pelleted by centrifuge at 2000g for 5 minutes at 4°C. For nuclei isolation, the pellet was resuspended using Cytoplasmic Lysis Buffer and incubate for 10 minutes on ice, pelleted at 600 × g for 5 minutes at 4°C, and resuspended in Nuclei Lysis Buffer. Chromatin was enzymatically digested with Micrococcal Nuclease (0.5 µl per 4 × 10⁶ cells) at 37°C for 20 minutes to produce fragments ranging from 150 to 900 base pairs, as confirmed by 1% agarose gel electrophoresis. The digestion process was halted with 10 mM EDTA. Nuclei were lysed via sonication (three 20-second pulses), and the lysates were clarified by centrifugation at 9,400 × g for 10 minutes at 4°C. For immunoprecipitation, 5 µg of chromatin was diluted in 1X ChIP Buffer and incubated overnight at 4°C with rotation using the following antibodies: 5 µl anti-PPARγ rabbit monoclonal antibody (#2443, CST); 2 µg normal rabbit IgG as negative control (#2729, CST), and 10 µl anti-Histone H3 rabbit monoclonal antibody as positive control (#4620, CST). Immune complexes were captured with ChIP-Grade Protein G Magnetic Beads (30 µl) for 2 hours at 4°C and subsequently washed by a series of buffers, including a low salt buffer, high salt buffer and elution buffer. Elution of chromatin was placed at 65°C for 30 minutes with gentle vortexing. This was followed by cross-link reversal, which involved incubation with 6 µl 5M NaCl and 2 µl Proteinase K at 65°C for 2 hours. DNA purification was achieved using spin columns with a DNA Binding Buffer, an ethanol-supplemented Wash Buffer, and a DNA Elution Buffer. The precipitated DNA was then analyzed via quantitative PCR using the SYBR Green Master Mix and primers specific to the *Slc25a34* promoter region (positions −327 to −59 bp relative to the transcription start site). The data were normalized to 1% input chromatin using the ΔΔCt method and expressed as a percentage of input, reported as the mean ± standard deviation (SEM) from three biological replicates.

### CUT & Tag seq and data analysis

CUT & Tag (Cleavage Under Targets & Tagmentation) strategy was conducted using Hyperactive Universal CUT & Tag Assay Kit for Illumina (TD903, Vazyme) according to the manufacturer’s instructions. In brief, differentiated wild type brown adipocytes were harvested and counted to ensure a cell viability of 90%. Pre-washed 1×10^5^ cells in 100 μl wash buffer in each sample were incubated with 10 μl pre-activated Concanavalin A-coated (ConA) beads for 10 min at room temperature with gentle mixing. After removal of the supernatant, the bead-bound cells were incubated with primary anti-PPARγ antibody at 1:50 (#2443, CST) in 50 μl antibody buffer at 4°C overnight. Following this, cells attached to beads were incubated with 50 μl of unconjugated secondary antibody at 1:100 dilution in Dig wash buffer for 1 hour at room temperature with rotation. After three times wash in 200 μl Dig-300 buffer, bead-bound cells were incubated with 0.04 μM pA/G-Tnp in Dig-300 buffer at room temperature for 1 hour, followed by additional three times wash with fresh Dig-300 buffer. Subsequently, the samples were treated with 10 μl TTBL in 40 μl Dig-300 buffer for DNA fragmentation through a 1-hour incubation at 37°C. Then the samples were treated with 5 μl Proteinase K, 100 μl of Buffer L/B, and 20 μl of DNA Extract Beads. The mixture was vortexed thoroughly and subsequently incubated at 55°C for 10 min, with two to three gentle mixing intervals during incubation to terminate DNA fragmentation and facilitate DNA extraction. After this, the beads were subjected to sequential washes with 200 μl WA buffer and 200 μl WB buffer. The fragmented DNA was eluted in 22 μl DNase-free water, of which 15 μl was used for library amplification following the manufacturer’s protocol. Amplified DNA was purified using 100 μl VAHTS DNA Clean Beads (#N411, Vazyme), re-eluted in 22 μl DNase-free water, and the normalized libraries of 5-10 nM final concentration were subjected to paired-end sequencing on the Illumina NovaSeq X Plus Series PE150 platform (Novogene) following standard cluster generation protocols.

### Oxygen consumption rate of cells

Cellular oxygen consumption rate (OCR) was assessed in fully differentiated brown adipocytes using a Mitocell (MT200) mixing chamber and a Model 782 oxygen Meter (Strathkelvin Instruments). Briefly, cells were scraped off and resuspended in 400 μL of cell culture medium to measure the basal OCR of the cells. Afterward, 5 mg/mL oligomycin A (S1478, Selleck) and 5 μM FCCP (HY-100410, MCE) were added to evaluate the uncoupled and maximum OCR, respectively. The OCR values were calculated using the analyze software (782 Oxygen System version 4.0).

### Stable isotope-labeled metabolome analysis

Stable isotope-labeled ^13^C6-glucose and ^13^C5-adenosine was utilized for metabolome analysis. Differentiated BAC were incubated with the glucose-free DMEM supplemented with 4.5g/L ^13^C6-glucose (CLM-1396-1, Cambridge Isotope Laboratories) or regular DMEM containing 20 μM ^13^C5-adenosine (CLM-3678-MG-PK, Cambridge Isotope Laboratories) for 24 hours. Metabolites were extracted post-incubation using a pre-cold solution of methanol and water (80:20,v/v) at 4°C with gentle rotation for half an hour, followed by centrifugation at 14000 × g for 10 minutes at 4°C. The supernatant was collected and transferred to the sample loading vials. Metabolites were isolated using an Ultimate 3000 UHPLC system (Thermo Scientific). The separation of compounds was performed with a ZIC-pHILIC column (2.1 mm i.d.×150mm, 5μm, Merck) at 30°C. The quantification of isotopomer distributions for key metabolites was achieved using isotopically resolved peaks.

### Structural model prediction

To generate structural models of AMP-SLC25A34 complex, the amino acid sequence of mouse SLC25A34 and canonical SMILES of AMP served as input for the prediction through a local installation of AlphaFold3. The parameter max_template_data was set to 2021-09-30. The other settings were default unless otherwise specified. The prediction output yielded a set of 5 ranked models representing the transport of AMP by SLC25A34. In addition to pLDDT and PAE, AlphaFold3 uses two further metrics, pTM and ipTM, to measure the accuracy of the predicted models. We used the top model, ranked by pLDDT, for the interaction analysis and figure creation by the software ChimeraX ^43^. The residues of SLC25A34, including CYS31, TYR93, GLN94, ASN98, ARG190, SER195, GLN198, ARG291, HIS295 and SER 299, were predicted to form polar interactions with AMP.

### Co-immunoprecipitation (Co-IP)

The coding sequence of mouse Slc25a34 and Slc25a20 was amplified from cDNA of BAT from C57BL/6 wild type mouse, and the coding sequence with Flag-tag were cloned into pcDNA3-vector to construct pcDNA3-Flag-Slc25a34 and pcDNA3-Flag-Slc25a20 expression plasmid respectively. Based on the predicted substrate-binding residues illustrated in Figure 2F, site-directed mutagenesis was conducted on the mouse Slc25a34 coding sequence. Specifically, the residues Cys31, Tyr93, Gln94, Asn98, Ser195, Gln198, His295, and Ser299 were individually substituted with alanine, while Arg190 and Arg291 were replaced with glutamic acid. These mutations were introduced simultaneously into the mouse Slc25a34 coding sequence. The mutant coding sequence with Flag-tag were synthesized by Jiangsu CoWin Biotech Co., Ltd and subsequently cloned into the pcDNA3 expression vector to construct pcDNA3-Flag-Slc25a34-mut expression plasmid. All constructs were verified through Sanger sequencing prior to subsequent experiments. HEK-293T cells were transfected with pcDNA3-vector, pcDNA3-Flag-Slc25a34, pcDNA3-Flag-Slc25a34-mut or pcDNA3-Flag-Slc25a20 using PEI and harvested 48 hours post-transfection. Cells were lysed in RIPA buffer (150 mM NaCl, 1% NP-40, 50 mM PH 8.0 Tris-HCl, 25 mM NaF, 2mM Na3VO4, 0.5% sodium deoxycholate, cocktail and 1mM PMSF) on ice incubation for 10 minutes. Lysates were then centrifuged at 13,000 rpm (4°C) for 10 minutes, and the supernatant was collected. For AMP detection, cleared cell lysates were incubated with pre-equilibrated FLAG antibody-conjugated beads (A2220, Sigma) for 6 hours at 4°C with rotation, followed by 5 washes with wash buffer (50 mM Tris-HCl, 150 mM NaCl, 1 mM MgCl2, 0.5% NP-40) and elution in 60 μL AMP buffer for AMP content measurement. For Slc25a34 protein detection, cleared cell lysates were incubated with AMP conjugated beads (A1271, Sigma) and unconjugated beads. The eluate from the beads was analyzed by western blotting using an anti-FLAG antibody.

### Mitochondria purification

Mitochondria of BAC was purified by immunoprecipitation (IP) following the procedures from a prior study ^44^. Immortalized brown preadipocytes stably expressing 3 × FLAG-EGFP-OMP25 were cultured and differentiated in 15 cm dishes. For IP, Protein A/G magnetic beads were pre-blocked with 3% BSA in PBS, then incubated overnight with an anti-FLAG antibody at 4°C. Prior to mitochondrial isolation, cells were washed once with 10 mL ice-cold PBS and followed twice with 10 mL ice-cold KPBS (136 mM KCl, 10 mM KH₂PO₄, pH 7.25). Cells were scraped using a cell scraper in 1 mL KPBS and homogenized on ice with a pre-chilled Dounce homogenizer for 20 strokes. The homogenate was transferred to a new 1.5 mL tube and centrifuged at 1,000 × g for 2 minutes at 4°C to pellet nuclei and unbroken cells. The mitochondrial-containing supernatant was carefully aspirated and added to the anti-FLAG antibody-bound beads, followed by incubation for 30 minutes at 4°C with continuous rotation. Then, beads complexes were magnetically captured. The unbound cytoplasmic fraction was centrifuged three times at 14,000 × g for 10 minutes at 4°C, and the final supernatant was stored at -80°C for subsequent analyses. Beads were subjected to three stringent washes with 1 mL KPBS to remove the unspecific binding, then rapidly resuspended in 1 mL cold KPBS. For quality control, 50 μL of the beads suspension was reserved for western blot analysis, and 25 μL of the beads suspension was resuspended in 50 μL KPBS and visualized via fluorescence microscopy to assess mitochondrial extraction efficiency. The remaining beads suspension was collected using a magnetic strip, and the supernatant was aspirated prior to metabolite analysis.

### FITC-labelled AMP treatment and visualization

FITC-conjugated AMP (adenosine monophosphate) was synthesized by Nanjing TGpeptide Biotechnology Co.,Ltd (TG-LG-22899). The FITC moiety was covalently conjugated to the exocyclic amino group at the C6 position of the adenine base of AMP, corresponding to the N⁶-amino site of adenine, while preserving the ribose and 5′-monophosphate functionalities. AMP-FITC was acquired as a powder. Molecular weight (736.60 Da) was confirmed by mass spectrometry. Purity (>95%; determined as 97.07%) was assessed by RP-HPLC using a C18 column (4.6 × 250 mm, 5 μm) with a trifluoroacetic acid/acetonitrile-water gradient. Structural identity and molecular mass were further validated by ESI-MS, showing a dominant peak matching the expected mass. On day 3 of BAC cell differentiation, the culture medium was supplemented with FITC-AMP at a final concentration of 5 μg/mL. After 5 hours of incubation at 37°C under 5% CO₂, cells were gently washed twice with pre-warmed DMEM and then the cells were stained with MitoTracker probe as previous described. Following staining, cells were rinsed twice with PBS to remove excess dye. Cells were treated with ice-cold 100% methanol for 15 minutes at 37°C, then washed three times with PBS. Fixed samples were mounted using Antifade Mounting Medium with DAPI (P0131-5ml, Beyotime). Imaging was performed using a fluorescence microscope (Nikon, TS2). Colocalization of MitoTracker and FITC were quantified by using ImageJ (v.1.53) and Colocalization Finder (v.1).

### Proteoliposome-based uptake assay

To prepare SLC25A34 proteoliposomes, a mixture of POPC and 18:1 Cardiolipin (Avanti Polar Lipids) at 20:1 (w/w) was dissolved in chloroform and subjected to overnight vacuum evaporation. The lipid film was hydrated in internal buffer (20 mM HEPES pH7.5, 150 mM NaCl, 10 mM AMP) and diluted to 10 mg/ml. After three cycles of freeze-thaw in liquid nitrogen to internalize the unlabeled AMP, the liposomes were extruded for 21 times through a 400 nm polycarbonate membrane with Avanti Mini-Extruder and destabilized with 0.45% Triton X-100. Purified SLC25A34 in DDM:CHS was added to the lipids at a protein:lipid ratio of 1:80 (w/w) and incubated at 4°C for 20 min. The detergent was removed by adding four sequential batches of Bio-Beads SM-2 (Bio-rad) under constant rotation at 4°C — three batches at 40 mg/ml for 20 min each, followed by a final batch at 360 mg/ml for 12 h. The Bio-Beads was removed and the supernatant was collected as SLC25A34 proteoliposomes. The external unlabeled AMP was removed through BeyoDesalt G-10 Midi desalting column (Beyotime), and the 0.5∼1 mL fraction was collected and concentrated to 100 mL using 100 kDa cutoff Amicon concentrator (Millipore). As a negative control, empty liposome reconstitution was performed in the similar manner, with buffer substituted for protein.

For the transport assay analyzed by fluorescence microscopy, proteoliposome at ∼4 mg/mL lipids were diluted with buffer (30 mM HEPES, 150 mM NaCl, pH 7.5) to 2 mg/mL and incubated with 10 μM FITC-AMP at 37 °C for 10 min. Transport reactions were terminated by adding ice-cold buffer. After removing the free substrate via Beyosalt G-25 spin column (Beyotime), the amount of AMP transported into the proteoliposomes was quantified by measuring the fluorescent intensity of the diluted proteoliposomes at 0.5 mg/mL using a microplate reader (TECAN) with excitation at 488 nm and emission at 525 nm at room temperature.

For isotope-based transport measurements, the SLC25A34 proteoliposomes (or control liposomes) were incubated with 1 mM ^13^C10-AMP at 37°C for 20 min, and subjected to BeyoDesalt G-10 Midi desalting column (Beyotime) to remove the unincorporated ^13^C10-AMP. Fraction of 0.5∼1 mL was collected and the liposomes were pelleted by centrifugation at 80,000 rpm for 10 min at 4°C and resuspended in 50% acetonitrile for following LC-MS/MS quantification.

### AMP content measurement

AMP levels in mitochondria and cytosol were quantified using a colorimetric assay kit (ab273275, Abcam) following the manufacturer’s protocol. Briefly, mitochondria and cytosol were homogenized in Assay Buffer XV and incubated on ice for 10min, and then centrifuged at 10000g, 4℃ for 10 min. Clear supernatants were transferred to a new tube and placed on ice.10 μl of sample diluted in 40 μl Assay Buffer XV was loaded into a 96-well plate and 50 μl of reaction mix containing 42 μl Assay Buffer XV, 2 μl Development Enzyme Mix III, 2 μl Developer V, 2 μl AMP Substrate Mix and 2 μl OxiRed Probe was added to samples, respectively. After incubation at 37°C for 60 min, absorbance was measured at 570 nm using a microplate reader. AMP concentrations were calculated by interpolating corrected sample values onto the standard curve and adjusted according to protein concentration.

### ADP and ATP content measurement

Mitochondrial and cytosolic ADP and ATP levels were quantified using the ADP/ATP Ratio Assay Kit (A552, DOJINDO) via luminescence detection, with ADP and ATP measured sequentially. Briefly, 10 μL of each sample was loaded into a white 96-well microplate. 90 μL of ATP working solution, prepared by combining ATP Substrate, Assay Buffer, and ATP Enzyme Solution, was added to each well. The plate was incubated at 25°C for 10 minutes to stabilize luminescence signals, which were then measured using a microplate reader; these values were quantified as ATP levels. Next, freshly prepared ADP working solution (premixed ADP Substrate and ADP Enzyme working solutions) was added to convert ADP to ATP. Following an 8-minute incubation at 25°C, luminescence intensity was measured. A background signal was acquired immediately before adding the ADP working solution, and ADP levels were calculated after normalizing to this background. All measurements were performed in triplicate for each experimental condition to ensure accuracy.

### Lactate content and PFK enzyme activity measurement

Lactate levels were quantified using the CheKine™ Micro Lactate Assay Kit (KTB1100, Abbkine). The plasma and the supernatant of cell culture medium were diluted in assay buffer as 1:40. A working reagent of 55 μl mixture comprising 31 μl Lactate Assay Buffer, 8 μl lactate dehydrogenase cofactor, 5 μl WST-8, 1 μl Enhancer and 10 μl Lactate Dehydrogenase was freshly prepared. 50 μl of samples were mixed with 50 μl working reagent in a 96-well plate and incubated at 37°C for 30 minutes. Then, the absorbance value at 450 nm was measured using a microplate reader. PFK activity was determined using the CheKine™Micro 6-Phosphofructokinase Activity Assay Kit (KTB1124, Abbkine) according to the manufacturer’s instructions. Cytosolic fractions of cells were obtained as previous described and Cytosolic fractions of tissues were isolated using a mitochondria isolation kit (C3606, Beyotime) according to the manufacturer’s protocol, with final centrifugation at 12000g for 10 minutes at 4°C to obtain the supernatant as the cytosolic extract. Reaction mixtures including 10 μl sample, 10 μl Enzyme 1, 10 μl Enzyme 2 and 170 μl Substrate Mix were incubated in a 96-well UV-Transparent microplate. Absorbance at 340 nm was measured at 20 seconds for A1 and 10 minutes 20 seconds for A2 using a microplate reader. ΔA (A1-A2) was used to calculate PFK activity and normalize to protein content of tissue and the numbers of cell samples. The final PFK activity was adjusted by PFK protein expression.

### Statistical analysis

All statistical analyses were performed using GraphPad Prism 10. Statistical differences were evaluated via two-tailed unpaired Student’s t tests for comparisons between two groups or analysis of variance (ANOVA) and appropriate post hoc analyses for comparisons of more than two groups. Analysis of covariance (ANCOVA) was used for statistical analysis of the VO2 and energy expenditure data, which was performed at https://calrapp.org. We use body weight as a covariate. Significance thresholds were established as follows: *P-value<0.05; **P-value<0.01; ***P-value<0.001. Each panel’s data includes statistical methods and P values, detailed in the figure legend.

## Acknowledgments

We thank Dr. Jiandie Lin from the University of Michigan for the plasmids Dr. Junli Liu from Fudan University for the *Adrb3*-KO mice and the members of the Zhao lab and Zhang lab for their delicate guidance and advice on this study. In addition, we would like to thank Drs. Lifeng Yang and Junyao Wang from the Mass Spectrometry Technical Platform at the Shanghai Institute of Nutrition and Health, Chinese Academy of Sciences, as well as Dr. Meng Liu from the Mass Spectrometry Technical Platform at Shanghai Jiao Tong University School of Medicine, for their valuable technical assistance. We thank Core Facility of Basic Medical Sciences, Shanghai Jiao Tong University School of Medicine, for assistance with electron microscope analysis. This work was supported by the Shanghai Frontiers Science Center of Cellular Homeostasis and Human Diseases.

## Author contributions

X.Y.Z., K. Z. and H.S. conceived the project and designed the research. Y.L., X.Y. and J.Z. performed the majority of the studies. H.S. performed some cell and animal experiments. J.X., W.L., K.W., and F.C. performed some animal experiments. Y.L., H.S., K. Z. and X.Y.Z. analyzed the data and wrote the manuscript with input from other authors.

## Corresponding author

Correspondence to Xu-Yun Zhao, Kaihua Zhang and Hongyong Song.

## Data availability statement

The authors confirm that the datasets used and analyzed during the current study are available from the corresponding author upon reasonable request.

## Ethics statement

The experimental protocol was approved by the University Committee on the Use and Care of Animals at Shanghai Jiao Tong University (Reference Number: A2022-064). The research was prospectively reviewed and approved by the Ethics Committee of Shanghai Ninth People’s Hospital (Reference Number: 2018-86-T77), and informed consent was obtained from the participants. All the research was conducted in accordance with both the Declarations of Helsinki and Istanbul.

