## supplemental Figures for "A Cold-Responsive Mitochondrial Transporter Stimulates Brown Adipose Tissue Thermogenesis"

### Supplemental Information

#### Figures

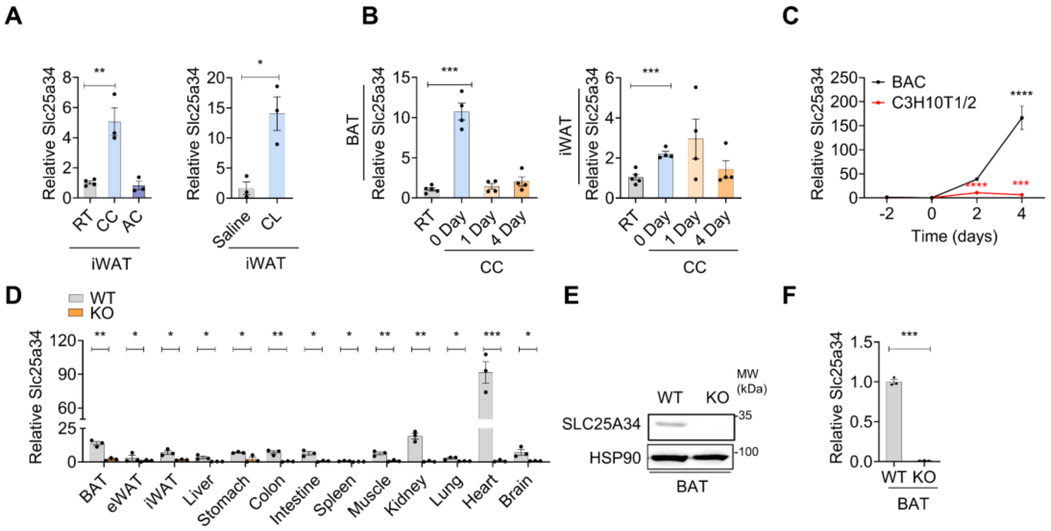


**Figure S1. Slc25a34 is induced by cold exposure and adipogenesis, and validation of Slc25a34-knockout mice**

(**A**) QPCR analysis of Slc25a34 expression in iWAT from mice treated in the room temperature (RT, n = 4), chronic cold (CC, n = 3), acute cold (AC, n = 3) and injected with CL316,243 (CL, 1 mg/kg, n = 3) or saline (Sal, n = 3) for 7 consecutive days. (**B**) QPCR analysis of Slc25a34 expression in BAT and iWAT from mice treated in the RT (n = 5), CC (n = 4) and reintroduction to RT for 1 day or 4 days following CC (n = 4). (**C**) QPCR analysis of Slc25a34 expression during BAC (n = 3) and C3H10T1/2 cells differentiation (n = 3). (**D**) QPCR analysis of Slc25a34 expression in various tissues from wild-type (WT) and Slc25a34-knockout (KO) mice (n = 3). (**E**) Western blot analysis of SLC25A34 protein in the BAT of WT and Slc25a34-KO mice. (**F**) QPCR analysis of Slc25a34 expression in immortalized WT and Slc25a34-KO brown adipocytes (n = 3). Data are presented as mean ± SEM. Statistical significance was determined by unpaired two-tailed Student's t-test in (D) and (F), one-way ANOVA multiple comparison test in (A) and (B) and two-way ANOVA multiple comparison test in (C). *P <0.05, **P <0.01, ***P <0.001. ns：not significant.

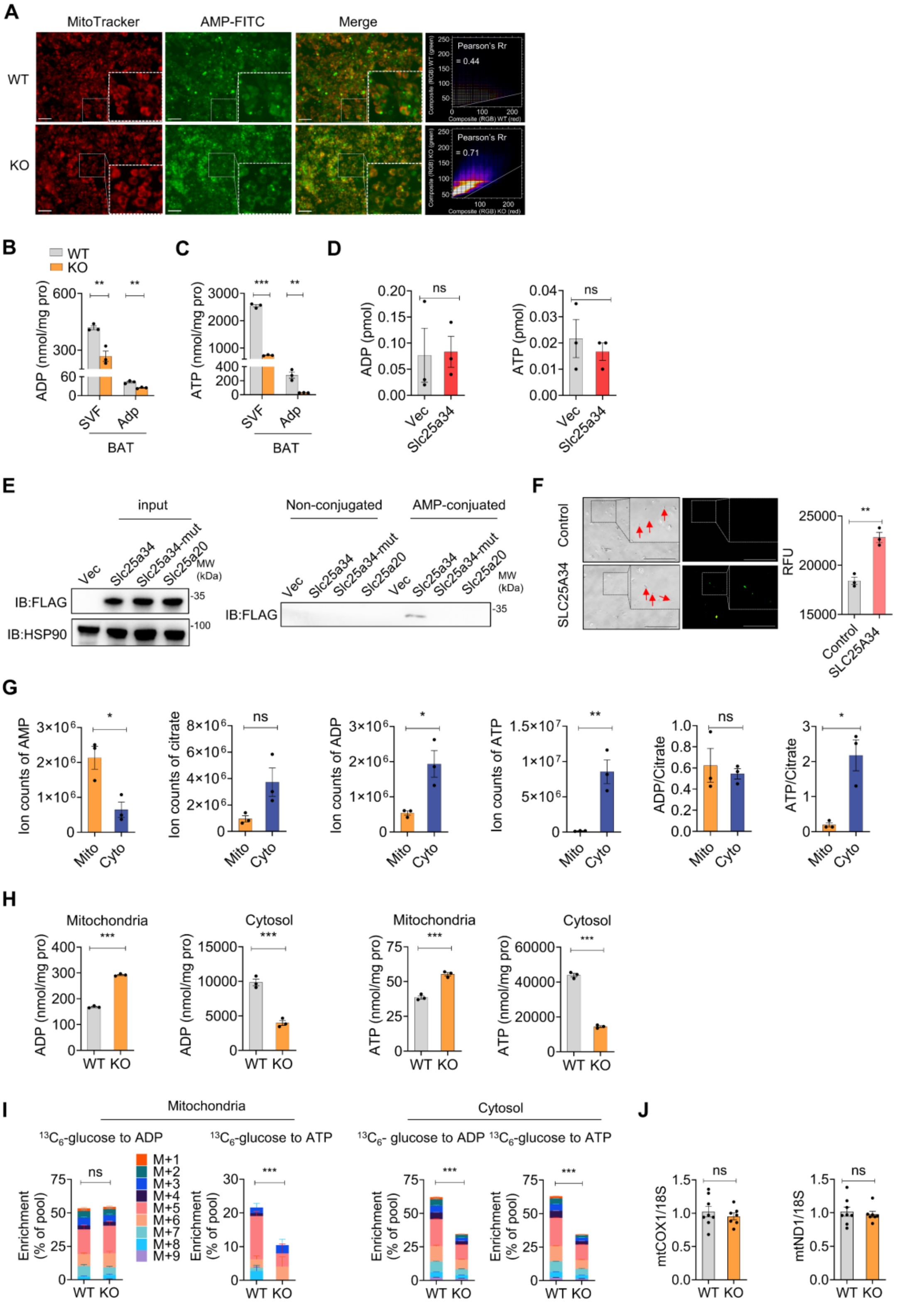


**Figure S2. Slc25a34 modulates adenine nucleotide metabolism**

(**A**) WT and *Slc25a34*-KO cells were treated with 10 μg/ml FITC-labeled AMP for 5 hours, then stained with Mitotracker. Subsequent fluorescence imaging was performed, with a scale bar of 100 μM. The colocalization of FITC and

Mitotracker signals was visualized and quantified using the Colocalization Finder program. (**B**) ADP concentration in stromal vascular fraction (SVF) and adipocyte fraction (Adp) of BAT from WT and Slc25a34-KO mice (n = 3). (**C**) ATP concentration in stromal vascular fraction (SVF) and adipocyte fraction (Adp) of BAT from WT and Slc25a34-KO mice (n = 3). (**D**) ADP (left) and ATP (right) content in immunoprecipitated beads from HEK 293T cells transfected with Flag-tagged Slc25a34 (n = 3). (**E**) HEK 293T cells transfected with Vec, Flag-Slc25a34, Flag-tagged Slc25a34-mut or Flag-tagged Slc25a20 (n = 3) were subjected to co-immunoprecipitation (Co-IP) using AMP-conjugated beads. The eluate was examined by western blot. The ratio of ADP and ATP to citrate levels in mitochondria and cytosol of brown adipocytes (n = 3). (**F**) SLC25A34 protein was reconstituted into proteoliposomes. FITC-labeled AMP was incubated with control proteoliposomes or those expressing SLC25A34 for 6 min, followed by acquisition of fluorescence images. Scale bar=100 μM. The relative fluorescence intensity was measured with a microplate reader (right). (**G**) The ion counts of Adenine nucleotide in mitochondria and cytosol of brown adipocytes. The ratio of ADP or ATP to citrate levels in mitochondria and cytosol of brown adipocytes (n = 3). (**H**)The content of ADP and ATP in mitochondria and cytosol in WT and Slc25a34-KO brown adipocytes (n = 3). (**I**) Mass isotopologue distribution analysis of ADP and ATP in mitochondria and cytosol of WT and Slc25a34-KO brown adipocytes following 24-hour incubation with $^{13}C_6$-glucose. (**J**) Relative mitochondrial DNA (mtDNA) copy number in BAT from Slc25a34-KO and WT mice. Data are presented as mean ± SEM. Statistical significance was determined by unpaired two-tailed Student's t-test in (B), (C), (D), (F), (G), (H), (I)and (J). *P <0.05, **P <0.01, ***P <0.001. ns：not significant.

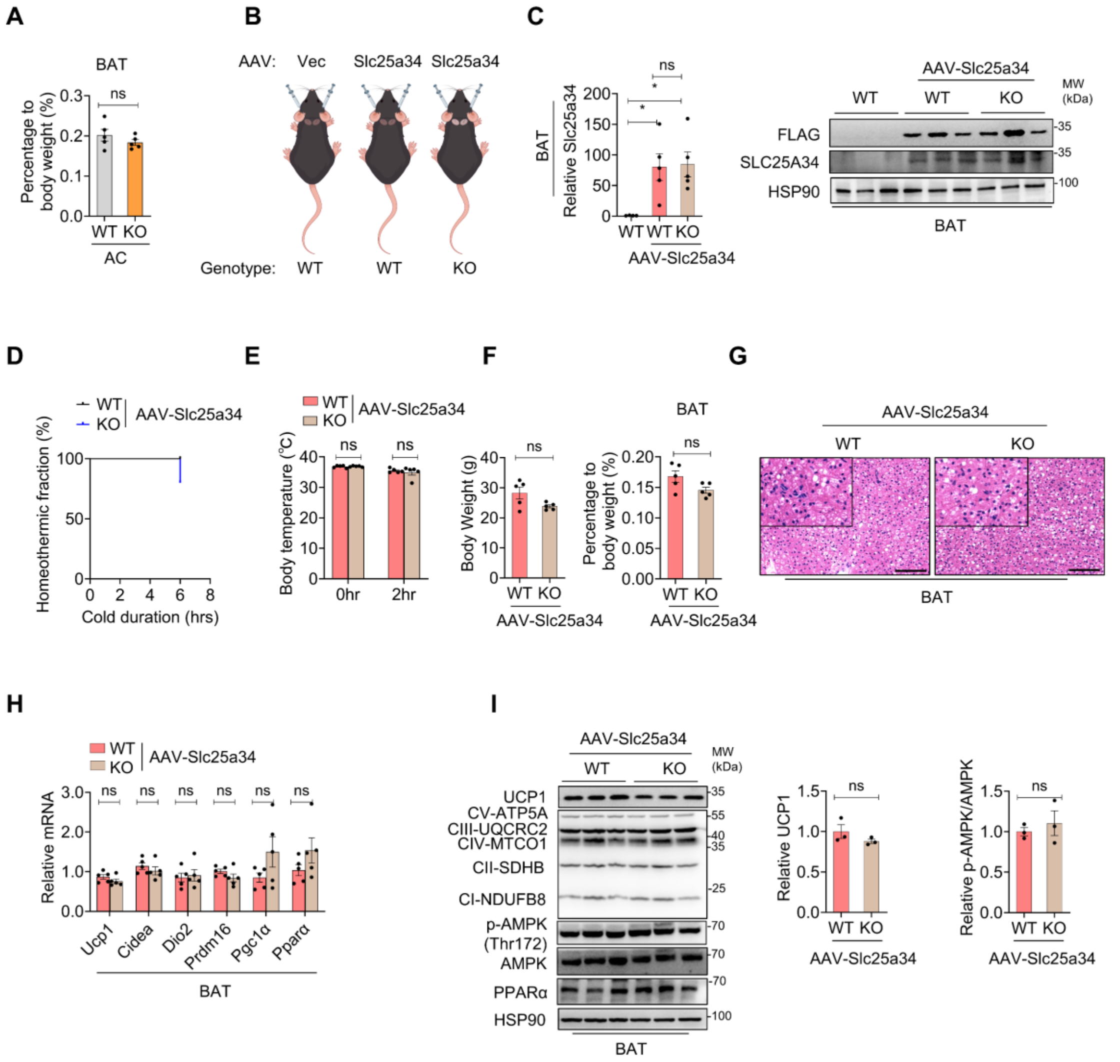


**Figure S3. BAT-specific SLC25A34 overexpression rescues BAT thermogenesis during cold exposure**

(**A**) Percentage of BAT weight to body weight of Slc25a34-KO mice (n = 5) and WT (n = 5) mice after acute cold exposure at 4°C. (**B**) Schematic showing the way of AAV injected in WT and Slc25a34-KO mice. (**C**) QPCR analysis of Slc25a34 expression in BAT of WT (n = 4), WT overexpression SLC25A34 (n = 5) and Slc25a34-KO overexpression SLC25A34 mice (n = 5) (left). The immunoblot analysis of Slc25a34 expression in BAT of WT (n = 3), WT overexpression SLC25A34 (n = 3) and Slc25a34-KO overexpression SLC25A34 mice (n = 3) (right). (**D**) The homeothermic fraction (rectal temperature>30°C) of WT (n = 5) or Slc25a34-KO (n = 5) overexpression SLC25A34 mice during 4°C acute cold exposure experiment. (**E**) Body temperature of WT (n = 5) or Slc25a34-KO (n = 5) overexpression SLC25A34 mice before and after 2 hours of cold exposure. (**F**) Body weight of WT (n = 5) or Slc25a34-KO (n = 5) overexpression SLC25A34 mice before and after cold exposure (left). Percentage of BAT weight to body weight of WT (n = 5) or Slc25a34-KO (n = 5) overexpression SLC25A34 mice before and after cold exposure (right). (**G**)Hematoxylin and eosin (H&E) staining of brown adipose tissue (BAT) from WT or Slc25a34-KO overexpression SLC25A34 mice subjected to acute cold exposure. Scale bar=100 μM. (**H**) QPCR analysis of mRNA expression of thermogenic genes in the BAT of WT (n = 5) or Slc25a34-KO (n = 5) overexpression SLC25A34 mice after acute cold exposure. (**I**) Western blot analysis of UCP1 and mitochondrial OXPHOS proteins in the BAT of WT or Slc25a34-KO overexpression SLC25A34 mice. The relative protein intensity of UCP1 in WT (n = 3) or Slc25a34-KO (n = 3) overexpression SLC25A34 mice (right). (**I**) Western blot analysis of phosphorylation of AMPK in WT (n = 3) or Slc25a34-KO (n = 3) overexpression SLC25A34 mice (right). Data are presented as mean ± SEM. Statistical significance was determined by unpaired two-tailed Student's t-test in (A), (E), (F), (H)and (I), one-way ANOVA multiple comparison test in (C), and Log-rank (Mantel-Cox) test in (D). *P <0.05, **P <0.01, ***P <0.001. ns：not significant.

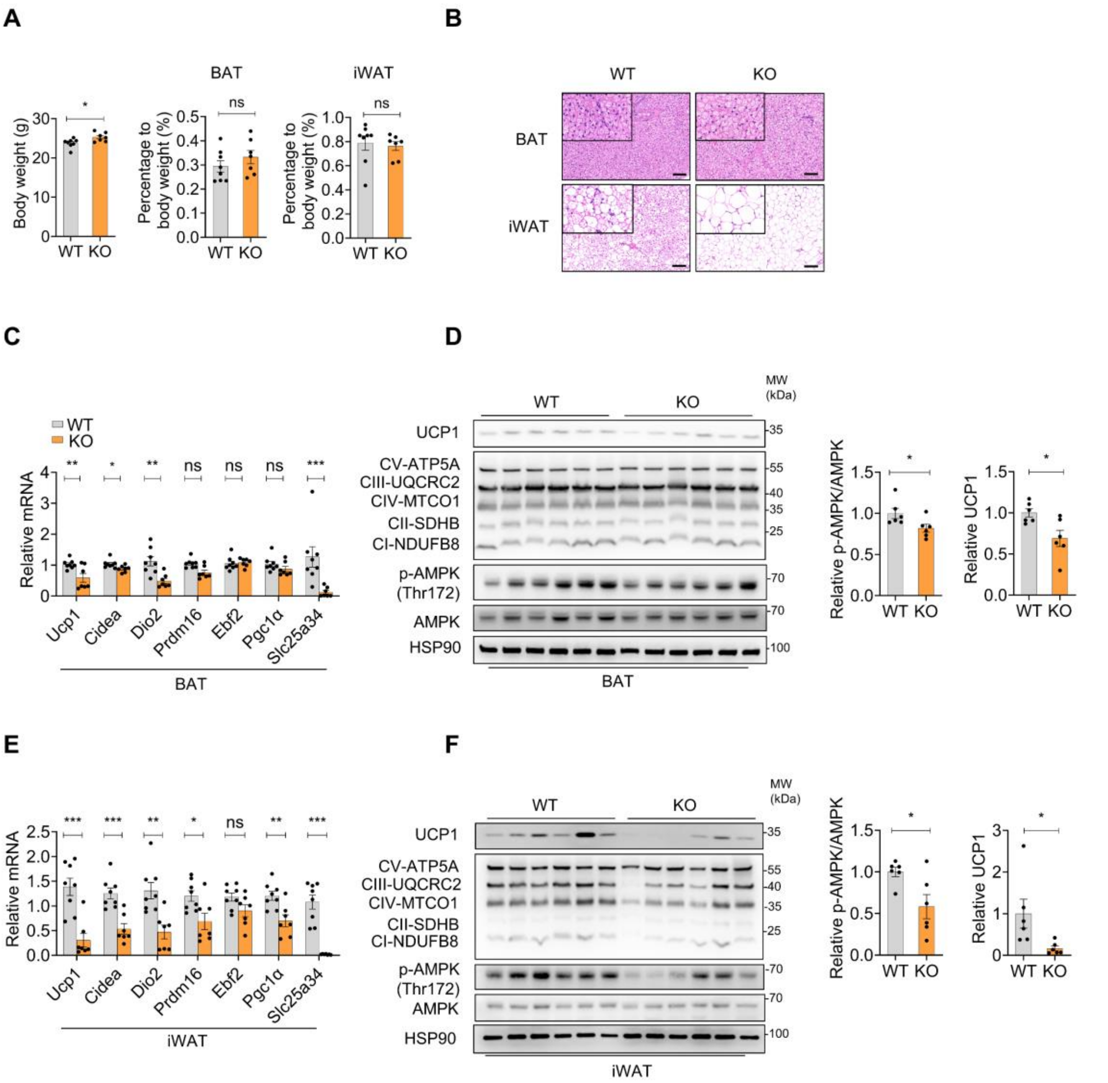


**Figure S4. Depletion of Slc25a34 inhibits CL316,243-induced thermogenic fat formation**

(**A**) Body weight of Slc25a34-KO (n = 7) and wild-type (WT, n = 8) mice following CL316,243 (CL) treatment. Percentage of BAT and iWAT weight to body weight of Slc25a34-KO (n = 7) and WT (n = 8) mice following CL316,243 (CL) treatment. (**B**) H&E staining of BAT and iWAT from mice after CL injection. Scale bar=100 μM. (**C-F**) Relative mRNA expression of thermogenic genes in BAT (C) and iWAT (E), and protein levels of UCP1, mitochondrial OXPHOS proteins, and phosphorylation of AMPK in BAT (D) and iWAT (F), from Slc25a34-KO (n = 7) and wild-type (WT, n = 8) mice following CL316,243 (CL) treatment. Data are presented as mean ± SEM. Statistical significance was determined by unpaired two-tailed Student's t-test in(A), (C), (D), (E)and (F). *P <0.05, **P <0.01, ***P <0.001. ns：not significant.

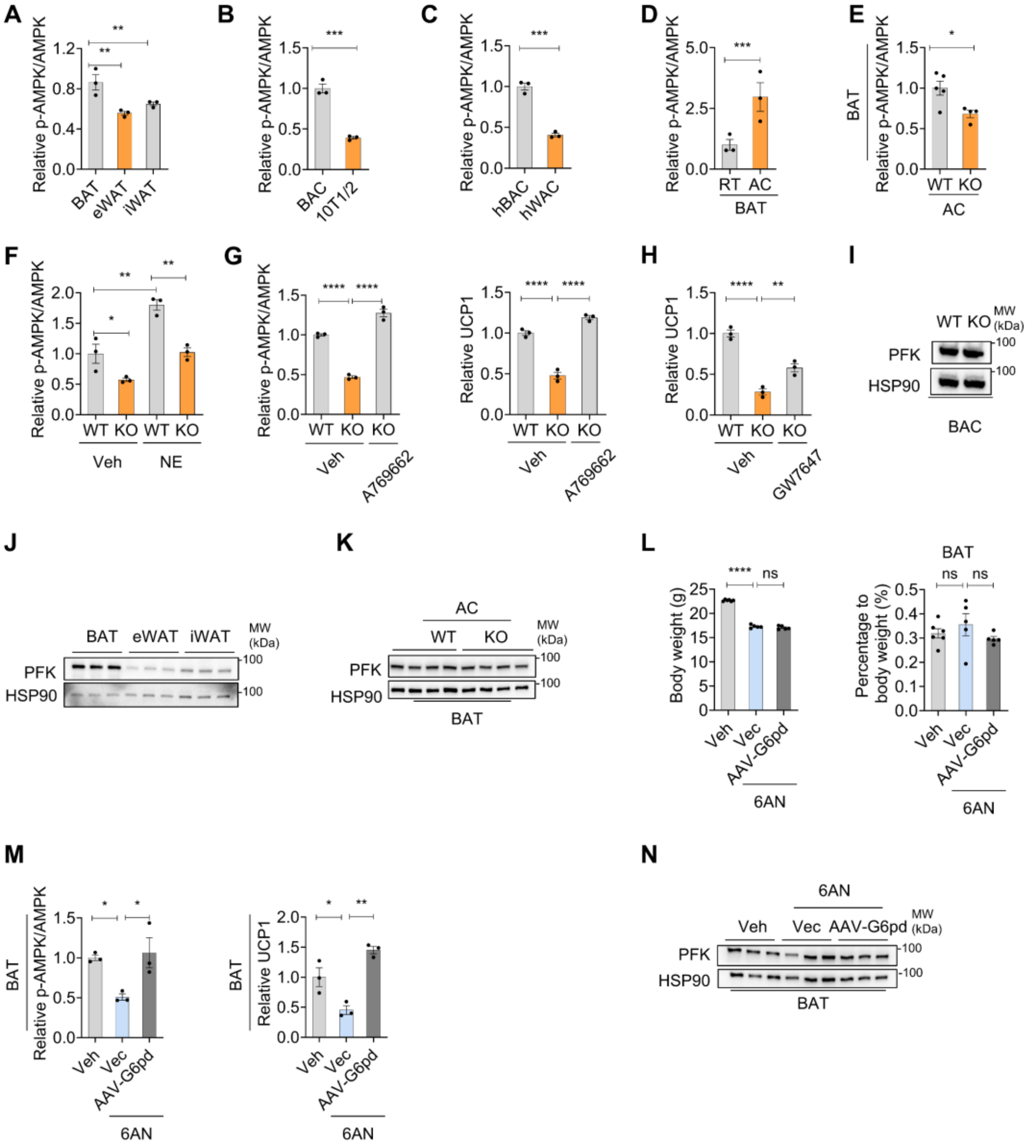


**Figure S5. SLC25A34 mediates the AMPK phosphorylation in BAT and BAC, and PFK protein expression regulated in glycolysis-TCA cycle flux**

**(**A)Phosphorylation of AMPK in different fat depots from WT mice housed at room temperature (RT). (**B**) Phosphorylation of AMPK in BAC and C3H10T1/2 (10T1/2) cells after differentiation. (**C**) Phosphorylation of AMPK in differentiated human brown (hBAC) and white adipocytes (hWAC). (**D**) Phosphorylation of AMPK in brown adipose tissue (BAT) from WT mice housed at RT or subjected to acute cold exposure at 4°C (AC). (**E**) Phosphorylation of AMPK in the differentiated WT and Slc25a34-KO BAC (n = 3) treated with vehicle (Veh) and NE (1 μM) for 6 hours. (**F**) Phosphorylation of AMPK in the BAT from WT and *Slc25a34*-KO mice subjected to acute cold exposure at 4°C (AC). (**G**) Phosphorylation of AMPK and relative UCP1 intensity in differentiated

WT and *Slc25a34*-KO BAC treated with Veh or A769662 (50 μM) during differentiation. (**H**) The relative UCP1 intensity in differentiated WT and *Slc25a34*-KO BAC treated with Veh or GW7647 (1 μM) during differentiation. (**I**) The PFK protein expression analyses were measured in differentiated WT and Slc25a34-knockout (KO) brown adipocytes. (**J**) The PFK protein expression analyses were measured in different fat depots from WT mice housed at room temperature (RT). (**K**) The PFK protein expression analyses were measured in brown adipose tissue (BAT) of WT and Slc25a34-KO mice following 4°C cold exposure (AC). (**L**) Body weight and percentage of BAT weight to body weight of Veh (n = 6), 6AN (n = 5), and G6pd overexpression mice subjected with 6AN (n = 5) for 5 days. (**M**)Phosphorylation of AMPK and relative UCP1 intensity in the BAT of mice injected with Veh, 6AN, and G6pd overexpression mice subjected with 6AN. (**N**) The PFK protein expression analyses was measured in the BAT of mice injected with Veh, 6AN, and G6pd overexpression mice subjected with 6AN. Data are presented as mean ± SEM. Statistical significance was determined by unpaired two-tailed Student's t-test in (B), (C), (D) and (E), and one-way ANOVA multiple comparison test in (A), (F), (G), (H) ,(L) and (M). *P <0.05, **P <0.01, ***P <0.001, **** P <0.0001. ns：not significant.

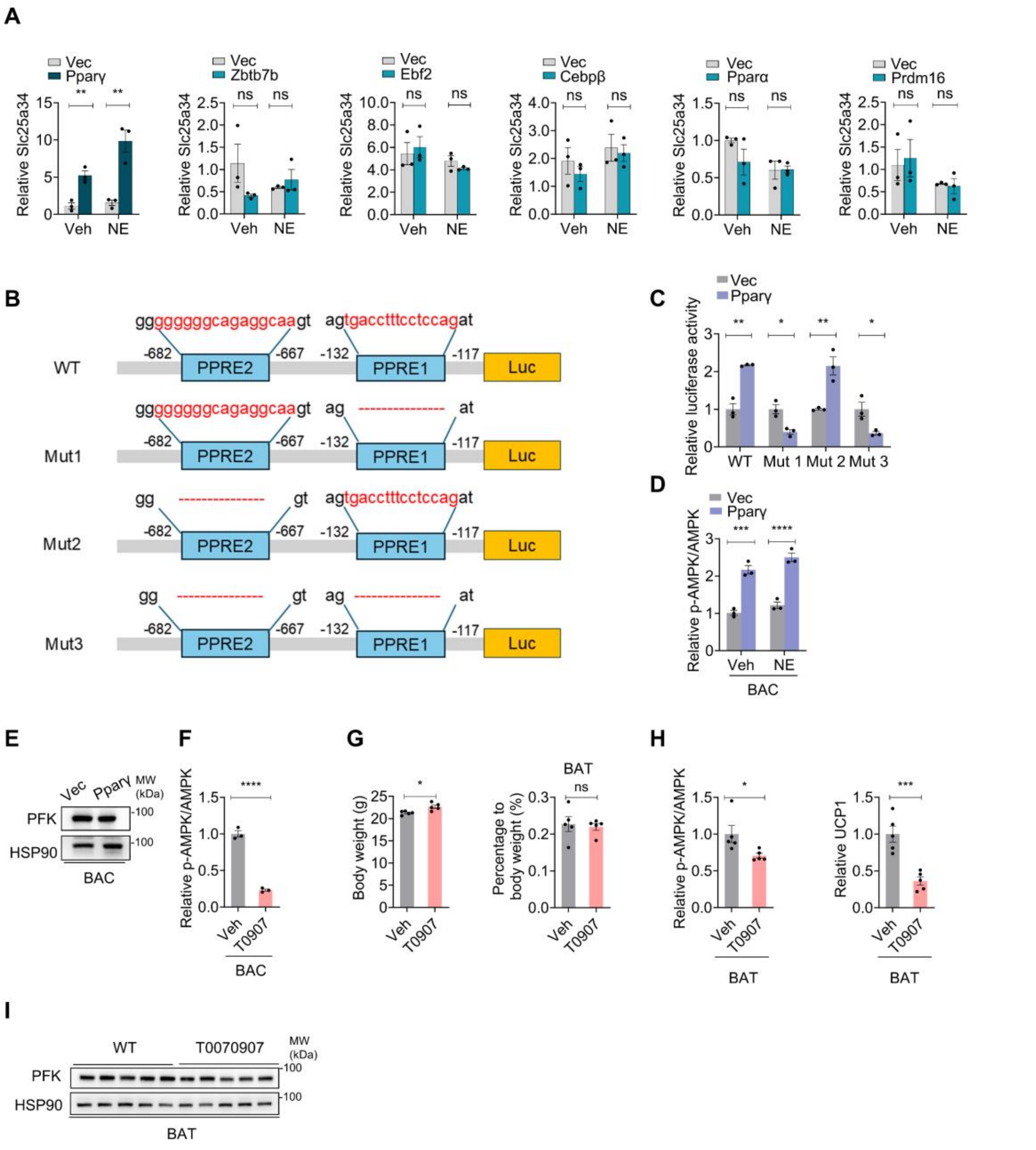


**Figure S6. A transcriptional factor screen identified that PPARγ transcriptionally regulates Slc25a34 expression**

(**A**) QPCR analysis of Slc25a34 expression in BAC overexpressing a panel of brown adipogenesis-associated transcription factors (n = 3). (**B**) Schematic of PPRE mutation sites in the pGL3-Slc25a34 promoter-luciferase vector. (**C**) Relative luciferase activity was measured in HEK293T cells (n = 3) transfected with the Slc25a34 wild-type or mutant promoter, in combination with empty vector (Vec) or PPARγ-expressing plasmids, respectively. (**D**) Phosphorylation of AMPK in differentiated brown adipocytes (BAC) overexpressing empty vector (Vec) and PPARγ, treated with vehicle and norepinephrine (NE, 1 μM) for 6 hours. (**E**) The PFK protein expression analyses were measured in Vec and PPARγ overexpressing BAC (n = 3). (**F**) Phosphorylation of AMPK in differentiated BAC treated with T0907 (10 μM) for 6 hours. (**G**) Body weight and percentage of BAT weight to body weight of Veh- (n = 5) and T0907-treated mice (n = 5) after 4°C cold exposure. (**H**)Phosphorylation of AMPK and relative UCP1 protein intensity in BAT of Veh- and T0907-treated mice after 4°C cold exposure. (**I**) PFK protein expression was measured in BAT of Veh- (n = 5) and T0907-(n = 5) treated mice after 4°C cold exposure. Data are presented as mean ± SEM. Statistical significance was determined by unpaired two-tailed Student's t-test in (A), (C), (D), (F), (G) and (H). *P <0.05, **P <0.01, ***P <0.001, **** P <0.0001. ns：not significant.

**Table S1. List of qPCR primers**

| Gene | Former primers (5’-3’) | Reverse primers (5’-3’) |
|---|---|---|
| mSlc25a34 | CTGCTGGCCTTCTCTACCAG | CCTGGTTGTTGGGTGAGGC |
| mUcp2 | CAGCGCCAGATGAGCTTTG | GGAAGCGGACCTTTACCACA |
| mUcp3 | CTGCACCGCCAGATGAGTTT | ATCATGGCTTGAAATCGGACC |
| mSlc25a51 | CAGGCTCAGATGTCACAGGTTA | TCAACATTGGGGGCCTCTTC |
| mSlc25a17 | TGCAGTGCTCCTGGAGATAAT | CTGTAGAAGAACGCTGACCTTT |
| mSlc25a1 | GAGGCATCGAAATCTGCATCA | GGATGGAGCCGTAGAGCAA |
| mSlc25a2 | GTGGCCTTTACAGGGGAACC | TTCCTGACAAACTGTTGGCAA |
| mSlc25a3 | GGCTCCATGAAGTATTATGCACT | AAACCACGAACGCCATCTTCT |
| mSlc25a4 | GAAGGGTCTCTACCAGGGTTT | AGTGTCATAGACTCCGAAGTAGG |
| mSlc25a5 | CAAGACAGCGGTAGCACCC | CGCAGTCTATGATGCCCTTGTA |
| mSlc25a10 | CGCAATCTACGAGACCATGC | GCCTCCAGTTAAACCACTGATG |
| mSlc25a11 | GTACCTCCCCTAAGTCTGTCAA | CTGCATCCGGTTCTTCACCAG |
| mSlc25a12 | ATGGCGGTCAAGGTGCATAC | AGTCATGTAATGCTCCCCGTC |
| mSlc25a13 | CAGCCCAACCCGAAAACTGT | CTGGAACGCCACCATAAACAA |
| mSlc25a14 | CCTGCGTTACTAAGACAGGCA | CTGACACTACCCCACAGATCA |
| mSlc25a15 | CAGCAGGTGGTCCGTAAAGTG | GGCATTTCACAAGCTCCGT |
| mSlc25a16 | CACGCCGAGACTTCTACTGG | ATGACGGTTGTGAGCTTGTAAT |
| mSlc25a18 | AGACCCGACTACAGAACCAG | AGCTGCCGCAAGAAGTCAT |
| mSlc25a19 | GTGGCGGGATCAGTGTCAG | TCCGTGGTATTTGGCATTGGG |
| mSlc25a20 | GACGAGCCGAAACCCATCAG | AGTCGGACCTTGACCGTGT |
| mSlc25a21 | GCCTCCAACGTCAGCTTACTG | GTTTTCACCACATCGAGAGGAT |
| mSlc25a22 | CATCGCTGGGCTAATCGGG | GCTCCCCTATACATGCCGAAG |
| mSlc25a23 | GCCTTGACCGGAACCAAGAT | CCATGCTGTGTAGGATTTTCTCT |
| mSlc25a24 | AGGCTTTCGGCAGATGGTAAA | CCTTCCTCGGTAAGCAACTTCT |
| mSlc25a25 | TGACCATCGACTGGAACGAGT | TCACCGACATCGAAGATCGTC |
| mSlc25a26 | GCACACGGACTCTACTTCACA | CTGCTTAACCACTTCAGAAGGAA |
| mSlc25a27 | TCTAACCACTTACGACACAGTGA | GCTTTACCGACCTTCCTTGTTT |
| mSlc25a28 | AGCATTGCGTGATGTACCCG | CCTGTTGCTGTGACGTTCA |
| mSlc25a29 | TGTGGCAGGTGTGATCGTG | GTGAGTCCCATTAGTGGCGA |
| mSlc25a30 | AGAGCCTGAAGCGGTTAGC | ACATCAGTCGGATTAGCAATAGC |
| mSlc25a32 | CACGTCCGGTACGAGAACC | TCCAAATGGTAGCCAAGCAAT |
| mSlc25a33 | CAGAAAGAGAACACGCTGCTT | GCAGTCGCGTCTTAATGACTT |
| mSlc25a35 | ATATCTGGGAAGCCCAATCTACA | CATGCCCTGATGCTTATACTGG |

| | | |
|---|---|---|
| mSlc25a36 | GCAACAAACCCCATTTGGCTT | GCCCCTATAAAATCCTCGCAG |
| mSlc25a37 | CCTACTCCACGATGCAGTAATG | AGTGAATTGACTGGAAGGGGATA |
| mSlc25a38 | TCCCCAGTGATCGAGAAGG | CTGGAAGAGGAGCGTGGAAC |
| mSlc25a39 | CTCGGCAACCAGCGAATTG | CCATTGCAGTATAGGAGGCACT |
| mSlc25a40 | TGGTGACTCCCCTGGATGTT | ATGTCCCACGGAAGTTTCCTG |
| mSlc25a41 | ATCGCCCCAGAGTATGCTATC | GCCGAGTCTTGAGTACCTCCA |
| mSlc25a42 | CCCTGGACCGGACCAAGAT | TGTATTCTTCGTGTGCGCTGA |
| mSlc25a43 | CAGGCATGGTTTCCACGATTG | GGAACAGCACCTAAAACAGTGAG |
| mSlc25a44 | GACGGTCTCCGAGGCTTCTA | TGGCTTGAAAGACAATGTGAGG |
| mSlc25a45 | GCACCCATTCGACACTGTAAA | CCCAGGACTGACTCATGGC |
| mSlc25a46 | GGACCAAGAGCCCTATGGAAA | CCTCCCTCGGTAAAGGTGTAAA |
| mSlc25a47 | GATTGGCAGTAACACCGGAAA | CCGTCTGGATTCTGACCTTCA |
| mSlc25a48 | CTGGAAGACTTTGTGGCAGG | GTTCGCATAGCCGACACCA |
| mSlc25a49 | TGGGCGATGTGGTTTTCTTGT | CCTGGCTAAACTGGGAACCT |
| mSlc25a50 | ACTGTGTTCAGGAGTCCTTGG | GCAGAACGAGCAATCATCTCTC |
| mPrdm16 | CGACTTTGGATGGGAGCAGA | GATTGGCATCCACGCAGAAC |
| mPparα | CGTCGGGATGTCACACAATG | GGCTTCGTGGATTCTCTTGG |
| mC/ebpα | GGCCAAGAAGTCGGTGGATA | GCGGTCATTGTCACTGGTCA |
| mC/ebpβ | CCAAGAAGACGGTGGACAAG | CACCTTCTTCTGCAGCCGCTC |
| mPparγ | CCGTAGAAGCCGTGCAAGAG | GGAGGCCAGCATCGTGTAGA |
| mZbtb7b | TCAGATGAGGATGCCATCGAT | CTTCCGCATGTGGATCTTCAG |
| mArrpp0 | GAAACTGCTGCCTCACATCCG | GCTGGCACAGTGACCTCACACG |
| mUcp1 | GGCATTCAGAGGCAAATCAGCT | CAATGAACACTGCCACACCTC |
| mDio2 | GATGCTCCCAATTCCAGTGT | TGAACCAAAGTTGACCACCA |
| mPgc1α | AGCCGTGACCACTGACAACGAG | GCTGCATGGTTCTGAGTGCTAAG |
| mPrdm16 | CGGAAGAGCGTGAGTACAAATG | TCCGTGAACACCTTGACACAGT |
| mCidea | GCAGCCTGCAGGAACTTATCAGC | GATCATGAAATGCGTGTTGTCC |
| mmtND1 | CCTATCACCCTTGCCATCAT | GAGGCTGTTGCTTGTGTGAC |
| mmtCox1 | CTACTATTCGGAGCCTGAGC | GCATGGGCAGTTACGATAAC |
| hSLC25A34 | GGTGCCTTGGAGACCATCTG | CCTCAGGGAGCCACTGTTG |
| hUCP1 | GCAGGGAAAGAAACAGCACC | CTTTCACGACCTCTGTGGGT |
| ChIP | ACAAGGAGGGGTGCCCTCC | CCTCTGGCTCAGACCAATC |
